# Transcriptomic module fingerprint reveals heterogeneity of whole blood transcriptome in type 1 diabetic patients

**DOI:** 10.1101/2025.11.19.689306

**Authors:** Felipe Leal Valentim, Océane Konza, Alexia Velasquez, Ahmed Saadawi, Nidhiben Patel, Roberta Lorenzon, Nicolas Tchitchek, Encarnita Mariotti-Ferrandiz, David Klatzmann, Adrien Six

## Abstract

**Background:** Type 1 diabetes (T1D) is a complex autoimmune disease resulting from β-cell destruction in the pancreas. Yet, no gene signature derived from whole blood samples has been established to differentiate T1D patients from healthy donors, likely because of the high heterogeneity of the underlying pathophysiological mechanisms.

**Methods:** We analysed whole blood transcriptomic profiles from 39 patients and 43 healthy donors, collected as part of our observational clinical trial, using a multi-step multivariate statistical analysis. This approach combined classical differential analysis, random forest and support vector machine classifications, gene set functional enrichment analysis to construct the most stable and reliable gene signature.

**Results:** Classical differential analysis did not separate clearly samples into healthy and T1D clusters, but rather spread samples into three clusters. On the contrary, we show that our approach, which combined molecular signatures independently constructed, adds robustness to the analysis without compromising the specificity. This efficiency was demonstrated by the clear separation of the samples according to their diagnosis group. Also, the functional annotation of the gene modules that we obtained was more associated with T1D-related pathways compared to the classical statistical analysis.

**Impacts:** These results emphasize that single differential analyses are not able to capture immune continuums involved in such a complex pathophysiological process. We hypothesize that T1D patients can have different molecular pathways involved in their pathology or/and can display unsynchronized -omics profiles. These findings call for further investigations to identify the molecular pathways involved in T1D patients, and for revising the nosology of autoimmune diseases.

## Introduction

Type 1 diabetes (T1D) is an autoimmune disease resulting from β-cell destruction in the pancreas (Lightfoot et al., 2012; Reynier et al., 2010; Tomita, 2017), which usually leads to complete insulin deficiency (American Diabetes Association, 2018; Petersmann et al., 2018). The molecular mechanisms underlying T1D are known to be related to antigen presentation (Felton et al., 2018; Mauvais et al., 2016), beta-cell autoimmunity (Knip et al., 2016; Petersmann et al., 2018), immune tolerance (Kuhn et al., 2016; Xu et al., 2013), T cell response (Burrack et al., 2017; Mallone et al., 2011; Roep, 2003), chemokine production (Collier et al., 2017), among other involved pathways (Felton et al., 2018; Polychronakos and Li, 2011; Roep and Peakman, 2012; Wållberg and Cooke, 2013). Understanding the pathogenic pathways that can explain β-cell destruction, as well as unveiling new regulatory components, is an open challenge towards the development of an immunotherapy for T1D (Boldison and Wong, 2016; Garyu et al., 2016; Kolb and von Herrath, 2017).

Genes that are related to a clinical outcome are collectively referred to as a “clinical molecular signature” (Bai et al., 2013; Tamez-Peña et al., 2018). Clinical molecular signatures have proved useful to predict prognosis, unveil the molecular mechanisms of disease, or be used as targets of novel treatments (Sim et al., 2017). These promises hold also for T1D molecular signatures that have been identified through GWAS and transcriptome analyses over the past years (Bakay et al., 2013; Nyaga et al., 2018; Størling and Pociot, 2017). Due to their potential, as well as recent advances in both high-throughput technologies with their respective bioinformatics methods, T1D molecular signatures have been incessantly studied during the last years (Eizirik et al., 2012; Ferreira et al., 2014; Kaizer et al., 2007; Reynier et al., 2010; Wang et al., 2008). However, due to a lack of analysis on the robustness of these molecular signatures across independent datasets (Park et al., 2012), and also due to differences among the experimental set-up and conditions, information from multiple molecular signatures are seldom if ever combined to explain molecular mechanisms underlying T1D.

One may hypothesize that a specific but broad, thus useful, clinical molecular signature for T1D would capture the intricate relationship between molecular mechanisms underlying β-cell destruction. Such molecular signatures can also be useful to help determine subgroups of T1D patients that can be distinguished due to a different level of disease activity and/or biochemical pathways involved. This remains unpractical for a single experiment due to cell type diversity, tissue specificity, and the heterogeneity in T1D (Chaussabel, 2015; Paterson and Petronis, 2000; Tritschler et al., 2017).

In this paper, we primarily aimed at establishing robust T1D signatures identified after an original data modelling scheme. To that end, we first identified a putative T1D molecular signature derived from our transcriptome expression dataset and assessed its specificity to cluster whole blood samples of T1D patients vs. healthy donors. We then harnessed information from the Molecular Signatures Database (MSigDB) (Liberzon et al., 2015) to identify literature-based or expert-based signatures that are also significantly enriched our expression set. At this step, we used a three-step analysis procedure to test each signature for enrichment, prediction, and clustering properties. All selected signatures have been combined with the original putative T1D molecular signature jointly analysed to identify transcriptomic modules and produce a T1D expression fingerprint, in a similar approach to that followed previously by Chaussabel et al.. We first use the transcriptomic modules to identify robust sample clusters. We then cross-analyse module expressions and clinical parameters to investigate sample cluster composition. Finally, we extend the current understanding of the molecular mechanisms underlying T1D by functionally annotating sets of modules that are specific to each diagnosis group and identified clusters.

## Results

### Description of the framework used for identification of a robust expression fingerprint of T1D in whole blood transcriptome data

In this work, we took advantage of the availability of a cohort of T1D and healthy volunteers recruited in the frame of our observational clinical trial Transimmunom (NCT02466217) as described in Lorenzon et al.. Whole blood from T1D patients (n=39) and healthy donors (n=43) was collected at two hospitals following a standardized protocol (Figure 1 and Materials and Methods). cDNA libraries were then prepared in batches to be sequenced either on HiSeq 4000 or NextSeq 500 machines. Sequencing reads were aligned to the Ensembl human genome and counted based on their overlap to known Ensembl genes. To estimate gene expression, resultant read counts were normalized and the batch effect was removed (see Materials and Methods). Differential expression (DE) analysis performed with four methods (DESeq2, wilcox, roc, t) was applied to this dataset to derive putative T1D gene signatures corresponding to the union of gene lists obtained from the four DE methods. Further, a previously developed method based on independent component analysis (ICA) was employed to generate additional potential signatures derived from the computed ICA sources computed from the transcriptome dataset (Pham et al., 2014).

**Figure 1.**
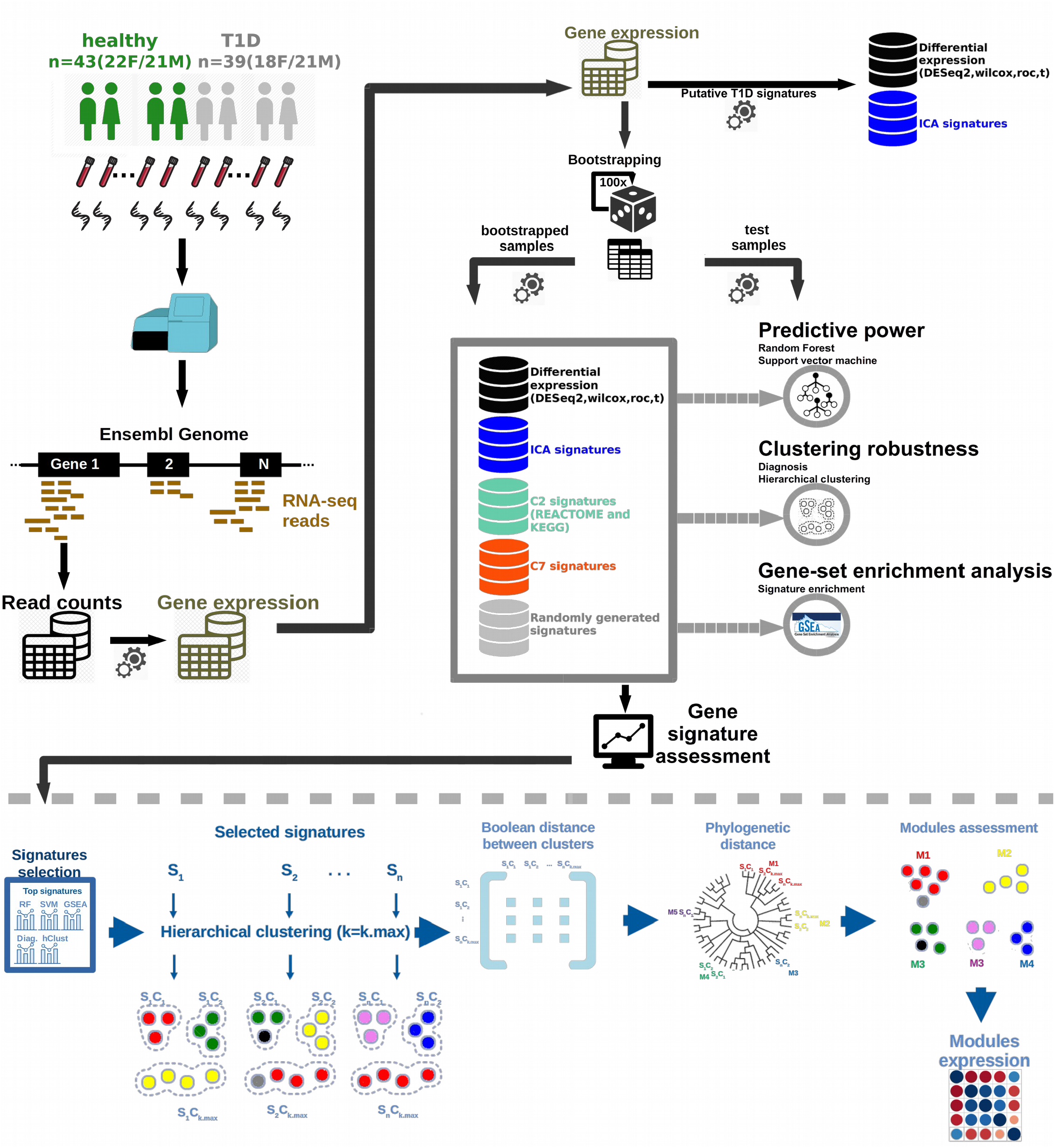
Schematic of the workflow for transcriptome analysis of T1D patients. Whole blood RNA from 43 healthy donors and 39 T1D patients were sequenced and the corresponding gene expressions were estimated. From this expression set, differential expression (DE) genes were obtained by four methods. Moreover, in addition to DE analysis, Independent component analysis (ICA) were also performed to generate putative molecular signatures. The Molecular Signatures Database (MSigDB) is an excellent public source of molecular signatures of multiple sources. From MsigDB resource, two gene set collections were considered: c2 (curated gene sets) and c7 (immunologic signatures gene sets). In addition, one randomly simulated gene set collection (Random) was also considered. For gene signature assessment, the initial sample set was bootstrapped 100 times by selecting 50% of samples for training the predictive models, while the remaining 50% of the samples were used for predictions assessment. Each of the signatures were evaluated in parallel by three groups of metrics: 1) the predictive power was measured by the accuracy of predictions obtained with either Random Forest or Support Vector Machine models, 2) the clustering robustness was measured with Calinski-Harabasz (CH) index calculated either from diagnostic or from independently computed clusters, 3) the ranking position of signature enrichment was calculated by gene-set enrichment analysis (GSEA). For producing a robust representation of T1D expression by combining all selected enriched signatures, a variation of transcriptomic module analysis is proposed. For this analysis, the genes in each molecular signatures are clustered by expressions, then the a distance matrix among all these gene clusters is constructed, from which a phyogenetic tree is constructed

We postulated that – with current high throughput technologies – a robust T1D molecular signature must contain information from several signatures to account for the roles of cell-type diversity, tissue specificity, heterogeneity, phenotypic plasticity (Ecker et al., 2018) and dataset robustness. However, a challenge in interpreting published clinical molecular signatures arises due to the limited robustness of independent datasets and differences among results (Sim et al., 2017). For this reason, a three-step analysis procedure was proposed here to assess previously published clinical molecular signatures. In addition to a “classical” gene-set enrichment analysis approach, we wanted to address two additional questions: “which signatures are better to classify patients?” and “which signatures produce more robust clusters of samples?”.

To answer the above-stated questions, we assess a signature by its 1) predictive power, 2) clustering robustness and 3) gene-set enrichment. The predictive power is given by the accuracy of predictions made with models constructed with the genes in a given signature, relying on two modelling approaches: Random Forest (RF) (Díaz-Uri-arte and Alvarez de Andrés, 2006) and Support Vector Machine (SVM) (Vanitha et al., 2015). The clustering robustness was computed for diagnosis-based clusters, but also for clusters obtained from hierarchical clustering (hclust). This second clustering approach was added to enable the identification of signatures that produce sub-clusters within the main diagnosis clusters. In both cases, the inter- and intra-distances of clusters were considered in the framework of the Calinski– Harabasz (CH) index (Caliñski and Harabasz, 1974) calculation as a proxy for clustering robustness. Finally, the gene-set enrichment analysis was used to rank the signatures according to the enrichment of gene-sets as a whole (Subramanian et al., 2005).

In addition to signatures generated by differential expression analysis (m=4) and ICA (m=2,726), gene sets from the Broad Institute molecular signature database (MsigDB) were also assessed, considering the gene set collections C7 (m=4,872) and C2 (m=4,738). For control pur poses, an additional 1,000 signatures were generated randomly. After assessing the ∼10,000 published molecular signatures, we selected enriched signatures according to metrics of predictive power, robustness and ranking from gene-set enrichment analysis. Aiming at producing a robust representation of T1D expression, we combined all selected enriched signatures by applying a variation of transcriptomic module analysis. Briefly, genes in each molecular signature are clustered according to their expression; then a distance matrix among all these gene clusters is computed and a circular hierarchical tree is constructed based on this matrix. This tree unveils gene clusters with similar gene composition which are then grouped into transcriptional modules. We finally produce an expression fingerprint representation by computing the average expression of genes in each module per sample, per diagnosis or per cluster. All steps of our modelling scheme are summarized in Figure 1.

### Putative DE-derived T1D molecular signature

We first performed a classical DE analysis on our expression set. Since it has been shown that sample size, effect size, and gene abundance greatly affect the results of all DE analysis methods (Jonsson et al., 2016; Quinn et al., 2018; Wesolowski et al., 2013), we used four methods to obtain the DE genes (DEG): a method based on negative binomial distribution model (DESeq2), the Standard AUC classifier (roc), Student’s t-test (t), and Wilcoxon rank sum test (wilcox). Considering these methods are complementary, we calculated the union of the four DEG lists obtained under stringent thresholds as presented in Table S1 (see Material and Methods for further information), thus generating our putative DE-derived T1D molecular signature.

On average, 152±82 DEGs (mean±sd) are detected by each method. The intersection between the four DEGs comprises 53 genes (Figure 2A) when the “DE union” signature comprises 269 genes. Here, a discrepancy between the number of up-(47) and down-regulated (22) genes in the “DE union” is observed. We also observed that a PCA on “DE union” genes (Figure 2B) shows a rather high separation on the x-axis moderately consistent with diagnosis labels. Since the optimal number of sample clusters observed from silhouette analysis was equal to k=3 (Figure 2C), we projected the k=3 clusters as computed by hclust onto the PCA space (Figure 2D): two prominent clusters can be found, one being 88% pure in healthy samples and another 78% pure in T1D. The third cluster is more heterogeneous showing 64% purity in healthy samples. To overview expression and sample clusters, a heatmap was generated (Figure 2E, see Material and Methods).

**Figure 2.**
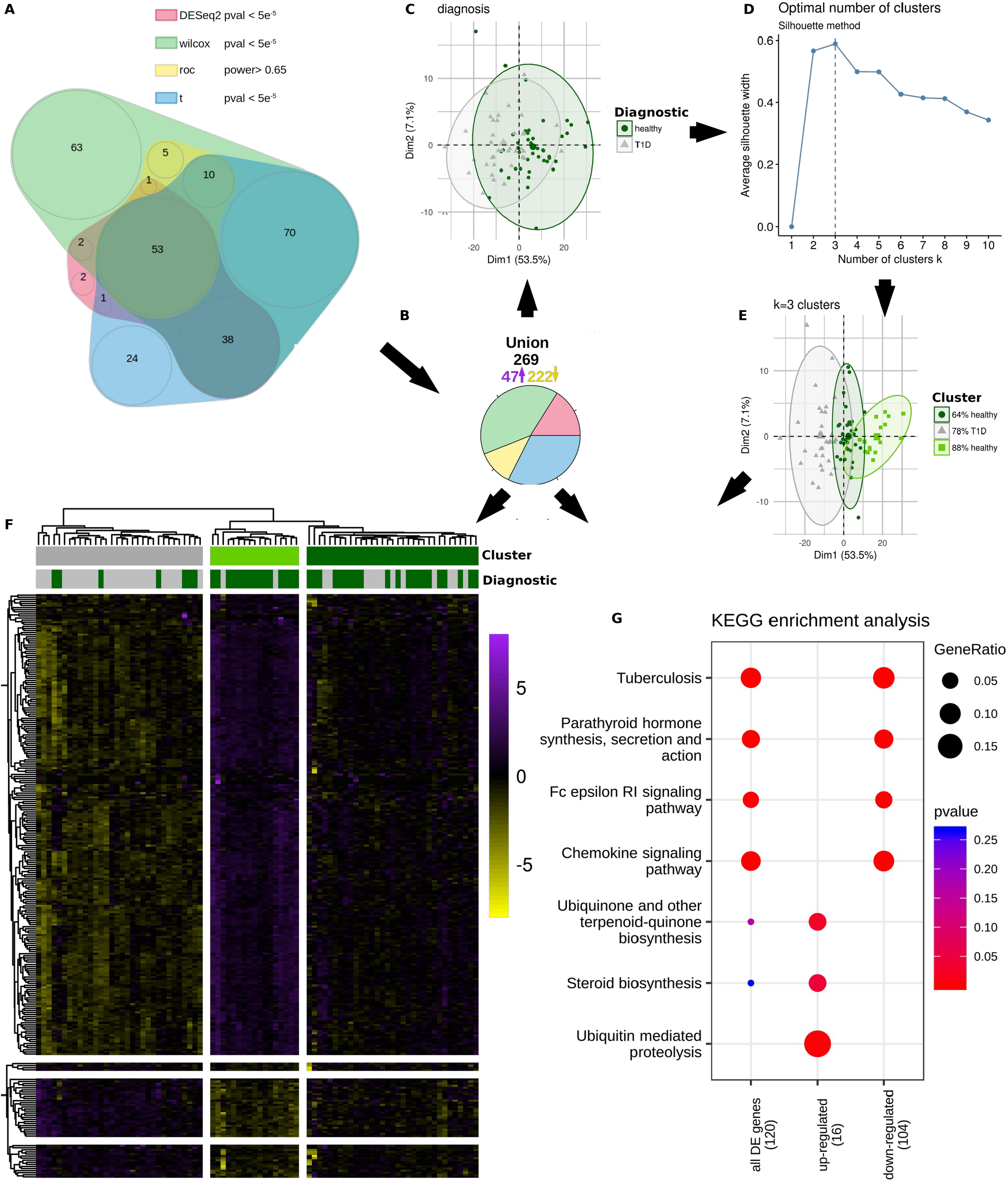
Description of the putative gene signature for T1D. Differential expression (DE) analysis performed with 4 methods (i.e. DESeq2, wilcox, roc, t) was applied to the expression set to derive a putative T1D gene signature. **A)** Overlap of the DE lists obtained from each DE method. The threshold used in each method is indicated. **B)** Schematic representation of union of DE lists obtained from each DE method. **C)** Principal component analysis on the putative T1D gene signature. D) Silhouette analysis for determining the number of sample clusters. The optimal cluster number is marked by dashed line. E) k=3 clusters delineated on PCA space. F) Heatmap presentation of the putative T1D gene signature. F) KEEG enrichment analysis of the putative T1D gene signature. Three groups were considered: all genes in the union of DE lists obtained from all DE methods, only the up-regulated genes within the union, and only the down-regulated genes.

Altogether, results show that our expression set captured specific features that are discriminatory of T1D diagnosis. Moreover, a functional analysis (Figure 2F) shows an overall enrichment of genes acting on auto-immune response (e.g. chemokine signalling pathway) and metabolic pathways (e.g. steroid biosynthesis and ubiquitin-mediated proteolysis). On T1D, we observe a down-regulation of genes related to auto-immune response concomitantly with an up-regulation of metabolic-related genes. We then aimed at improving the construction of gene signatures to better separate healthy volunteers and T1D patients by adding genes from independent molecular signatures. We thus implemented the proposed three-fold assessment to identify independently constructed molecular signatures that are enriched in our healthy or T1D transcriptome datasets.

A three-fold assessment identifies additional T1D-related signatures

The prediction of clinical status using gene expression profiles aims at classifying patients given a molecular signature. In the field of oncology, this approach helps to identify genes that are differentially expressed in tumours with different outcomes, so treatment strategies can then be tailored to groups of patients (von Herrath, 2005; Wei et al., 2004; Zarringhalam et al., 2018). In this work, we were primarily interested in obtaining the classification performance of models constructed with molecular signatures. The classification performance is used as a quantitative estimate of the signature predictive power. This strategy has already been implemented (Cha et al., 2016). Here, we wanted to identify molecular signatures with higher predictive power for diagnosis and thus retained the top ten signatures with the best predictive power determined by the accuracy of RF models or accuracy of SVM models (Table S2).

Overall, whilst one would expect the two metrics for prediction power to be highly correlated, only a medium correlation (≥0.5) is seen between accuracies from RF and SVM models. Strikingly, there is a better agreement (correlation ≥0.6) between clustering robustness (from diagnosis) and accuracy from SVM model (). Moreover, a bootstrapping procedure was implemented to explicitly account for heterogeneity, in such a way that each signature was evaluated on 100 different sub-datasets. A high (≥0.6) correlation among metrics on bootstrapped sub-datasets was seen () for clustering robustness metrics (mean correlation ± SD diagnosis = 0.66±0.69; robustness hclust = 0.64±0.19), a medium correlation (≥0.5) for gene-set enrichment rank analysis (GSEA rank = 0.50±0.16), and a low correlation (≤0.5) for predictive power (accuracy RF = 0.22±0.23; accuracy SVM = 0.32±0.27).

Analysis of each assessment metric averaged per signature collection consistently shows DE as the top scoring group (see boxplots in Figure 3A-E). However, few ICA signatures detected as outliers outperform the DE signatures for the metric of clustering robustness from hclust. This can be attributed to the nature of the ICA analysis, which detects signals from concise groups regardless of the sample labels (i.e. the diagnosis). Finally, but not least, c2 and c7 signatures perform significantly better than random signatures for all metrics. Based on these, we proceeded to evaluate signatures regardless of the collections they belong by taking the top ten signatures from each evaluated metric. These top signatures are hereafter referred to as the enriched signatures.

**Figure 3.**
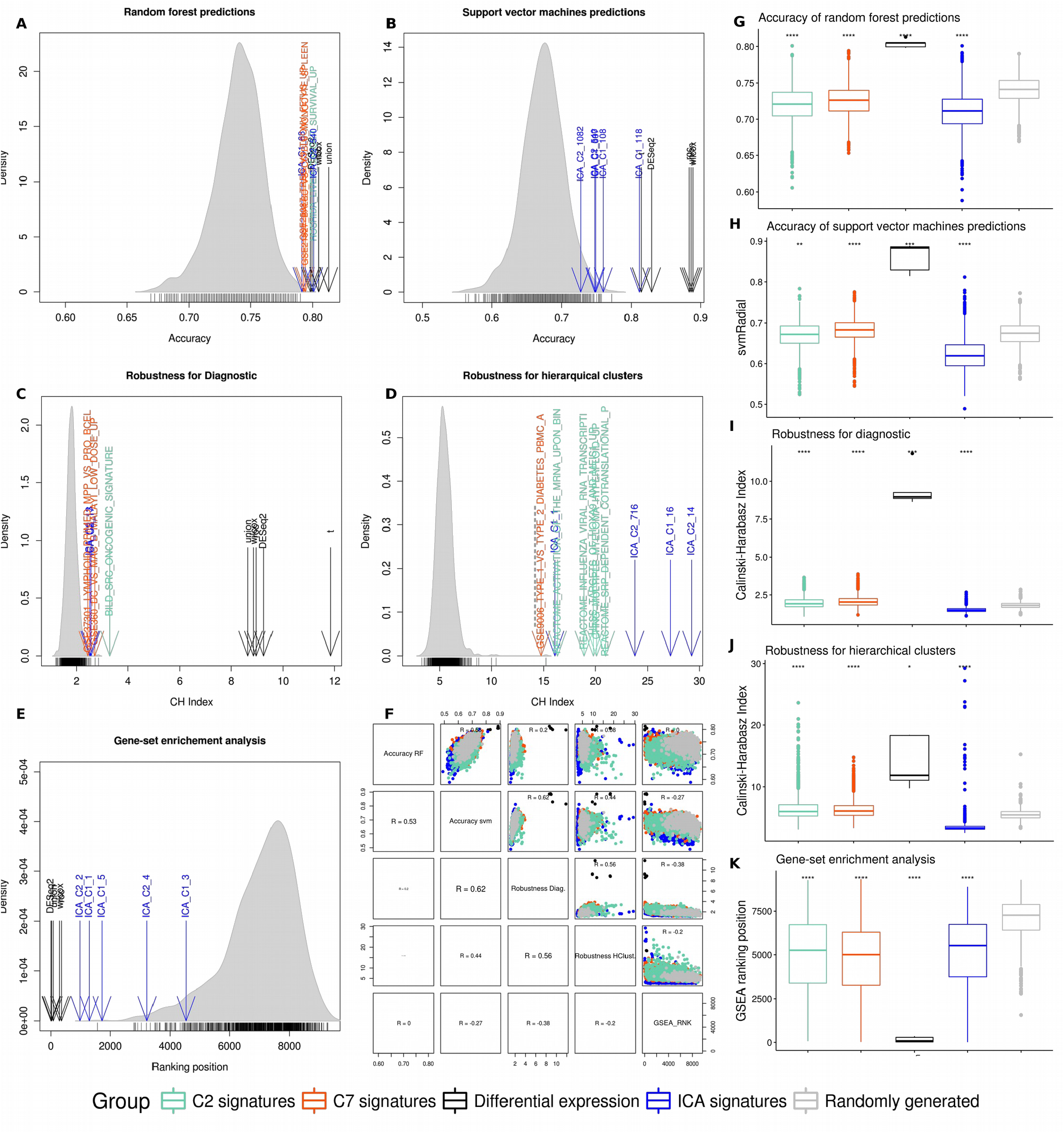
Quantitative assessment of gene signatures. The 1000 randomly-generated gene signatures were used to compute all histograms as well as to calculate p-value for the significance of enrichment (see Materials and Methods for further information). The values for the top 10 enriched signatures for each evaluated metric are represented by arrows. **A-B)** The predictive power was measured by the accuracy of predictions obtained with either (A) Random Forest (Accuracy of Random Forest)(A) or (B) Support Vector Machine (Accuracy of SVM) (B) models. **C-D)** The clustering robustness was measured with Calinski-Harabasz (CH) index calculated either from (C) diagnosis (C) (Robustness for Diagnosis) or from (D) independently-computed (D) hierarchical clusters (Robustness for hClust). **E)** Ranking position of signature enrichment was calculated by gene-set enrichment analysis (GSEA RNK). **F)** Correlation matrix Matrix of scatter plots between all pairwise combinations of metrics used to evaluate the signatures (i.e. Accuracy of Random Forest, Accuracy of SVM, Robustness for Diagnosis, Robustness for hClust and GSEA RNK). Pearson’s correlations are indicated. **G-K)** Assessment of signatures grouped by collection set (DE, ICA, C2, C7 and random). All t test comparisons were made against the reference collection set Random. For boxplots: (t test, ns: p > 0.05, *: p ≤ 0.05, **: p ≤ 0.01, ***: p ≤ 0.001, ****: p ≤ 0.0001)

From predictive power analysis, we primarily observed that all the DE signatures are enriched according to predictions from both RF and SVM models. Furthermore, we observed that two ICA signatures are enriched according to RF models, and five according to SVM models. One signature (HOSHIDA_LIVER_CANCER_SURVIVAL_UP) from c2 gene set collection, and two other signatures (GSE25087_TREG_VS_TCONV_FETUS_UP and GSE21927_BALBC_VS_C57BL6_MONO-CYTE_SPLEEN_DN) from c7 gene set collection, are enriched according to RF models. We then checked whether the signatures selected by their predictive power are overall enriched with T1D-related genes. For that, we ranked the genes according to variable importance (varImp) analysis (Figure S1 and Material and Methods). The top 20 genes - sorted by varImp - from each signature were further considered. Finally, we counted in how many enriched signatures each selected gene appears. On top of count list, MTRNR2L1 (ENSG00000256618) belongs to genetic locus previously associated to a “insulin-resistant obese cohort” (Nazarenko et al., 2017). The second on the count list, CCDC34 (ENSG00000109881) appears as up-regulated in replicating beta-cells (Klochendler et al., 2016). The third is the CLN3 (ENSG00000188603), which appears as DE on a recent study of the effects of T1D risk alleles on immune cell gene expression (Ram and Morahan, 2017). Still on the count list, we also find genes such as CYTH2, GCSAML, RIPK3, TUBD1, BCAR3, NDFIP2, RNASE1, LRRN3, REG4, SEMA6B, BTNL3, FAM153C, HLA-DQA1, LPAR3, RPS26 and SIGLEC11, all of which have been identified as genes with higher expression levels in mature β-cells compared to in vitro-differentiated insulin-positive β-like cells (Melton and Hrvatin, 2014) (Table S3).

Another tested hypothesis was that molecular signatures enriched in T1D-related genes will produce the most robust sample clusters. The Cal-iñski–Harabasz (CH) index was used as estimate of clustering robustness (see Material and Methods for further information). We first analysed clustering robustness (CH index) based on gene expressions grouped by diagnosis. Similarly to what has been done for predictive power analysis, we start by taking the top ten signatures (Table S2). Here, we primarily observed that all DE signatures are enriched. Furthermore, we also observed that two ICA signatures are also enriched. Interestingly, one signature (BILD_SRC_ONCO-GENIC_SIGNATURE) from c2 gene set collection, and two other signatures (GSE360_DC_VS_-MAC_B_MALAYI_LOW_DOSE_UP and GSE37301_LYMPHOID_PRIMED_MP-P_VS_PRO_BCELL_DN) from c7 gene set collection, appear as enriched here.

We then visually inspected plots of principal component analysis (PCA) constructed with the signatures. From PCA plots (Figure S2), we observed that the clustering robustness as measured by CH index is also effective for sample separation per diagnosis. This observation holds true for all the enriched signatures, but unsurprisingly most prominently for the DE signatures.

We also used clustering robustness definition to identify signatures that produce sub-clusters within the main diagnosis clusters. For this, instead of the original diagnosis label of each sample, we applied hierarchical clustering on expression levels of genes in each signature. An optimal number of k=4 clusters were produced (see Material and Methods for further information) and the CH index was computed over the obtained labels. Again, the top ten signatures were considered. Interestingly, we see the enrichment of a T1D signature (GSE9006_TYPE_1_VS_TYPE_2_DIA-BETES_PBMC_AT_DX_UP) from c7 gene set collection obtained from transcriptome analysis of peripheral blood mononuclear cells from children with diabetes (Kaizer et al., 2007). We proceeded with an investigation of the clusters obtained from expression of the genes in this T1D signature, and we find a very concise cluster of 22 samples with 77% purity in healthy samples (Figure S3). Moreover, we found that the c2 signature HESS_TAR-GETS_OF_HOXA9_AND_MEIS1_UP also produces prominent clusters, with sizes 33 (79% purity in healthy samples) and 23 (74% purity in T1D samples) respectively. Altogether, we reinforce the assumption that these signatures contain gene sets that concisely separate only a subset of samples due to heterogeneity.

Despite a variety of methods for gene-set enrichment analysis (Barry et al., 2005; Mathur et al., 2018; Tian et al., 2005), we relied on the popular GSEA (Subramanian et al., 2005) (see Material and Methods). This analysis was added to produce a rank for the signatures that account for all genes in the signature simultaneously, accounting for the possibility that a signature may contain only a few but highly discriminatory genes (such as expected from some gene-sets in the c7 immunologic signatures). Overall, we observed that on average DE signatures rank on top from GSEA analysis (p ≤ 0.0001), followed by c7 (p ≤ 0.0001), c2 (p ≤ 0.0001), ICA signatures (p ≤ 0.0001), respectively (Figure S4).

### Towards a T1D expression fingerprint

After identifying our putative DE-derived T1D signature and exploring ∼10,000 literature-based molecular signatures to select the top enriched ones, we aimed at producing a robust composite representation of T1D expression by combining all selected signatures presented on Table S2. Aware of the possible difficulty to interpret such a large number of signatures (34) and genes (3,793) as a whole (Ioannidis, 2005; Michiels et al., 2005; Santos Rego et al., 2013), we opted to reduce numbers by defining transcriptomic modules. Briefly, in a first step, we computed k gene clusters from each molecular signature (k estimated as the optimal number of clusters). In the second step, we calculated a distance matrix among all gene clusters using binary distance, then computing a distance tree. Each node of the distance tree represents one of the gene clusters derived from selected molecular signatures (Figure S5). ‘Transcriptional modules’ are then created by cutting neighbouring nodes in the binary distance tree. Finally, the average expression of genes in each ‘module’ is computed for each sample, cluster, and diagnosis condition (Figure S6). Table S4 shows statistics between module expressions and diagnosis (see Material and Methods for further information).

To focus on contrasting, robust and informative modules, we applied a multi-criteria filter, retaining modules whose a) expressions correlate with diagnosis (chi-square test pval < 0.05), and b) have significant mean difference (t-test pval < 0.01) between the T1D and healthy groups, and 3) appear among the top ten on the list ranked by variable importance analyses from a random forest model (Figure 4A).

**Figure 4.**
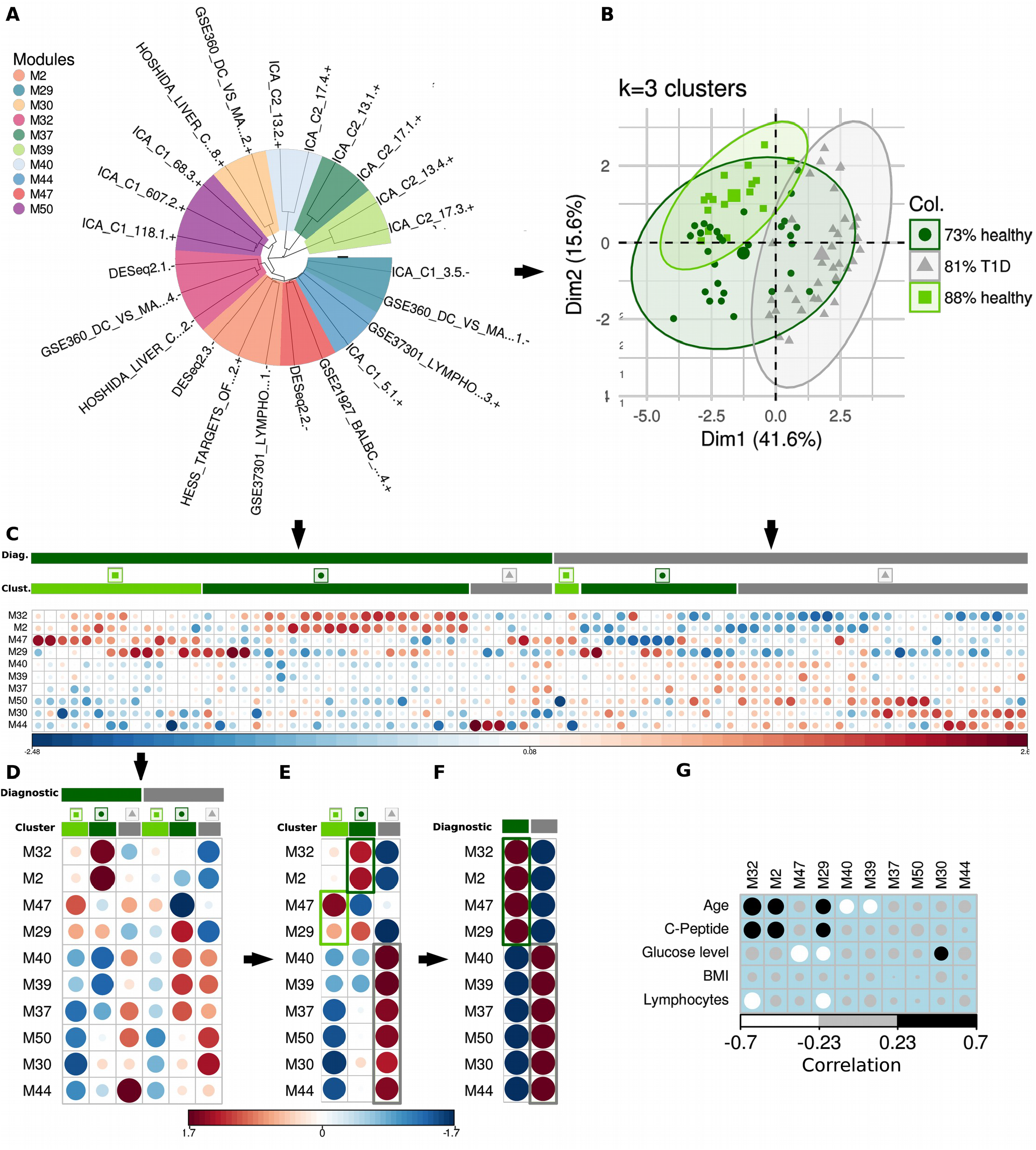
Transcriptomic module expressions for T1D. A) Phylogenetic representation of distance tree of selected modules. **B)** Principal component analysis on transcriptomic module expressions in which k=3 clusters are delineated. **C)** Transcriptomic module expressions per sample. **D)** Transcriptomic module expressions per clusters and diagnosis. **E)** Transcriptomic module expressions per cluster regardless of the diagnosis. **F)** Transcriptomic module expressions per diagnosis regardless of the cluster. **G)** Correlation between transcriptomic module expressions (per sample) and clinical parameters. We hereafter regard the modules M47 and M29 as exclusive for the cluster “88% healthy”, the modules M2 and M32 as exclusive for the cluster “73% healthy”, and the modules M40,M39, M37, M50, M30 and M44 as exclusive for the cluster “81% T1D”. Similarly, we hereafter consider the modules M2, M32, M47, and M29 as exclusive for healthy samples, and M40,M39, M37, M50, M30 and M44 as exclusive for the cluster T1D samples. These are delineated in the plots.

We then plotted the PCA from module expressions and delineated l = 3 sample clusters (Figure 4B) similar to our initial cluster analysis based on the DE signature only. Two of these three clusters obtained after hierarchical clustering on module expressions are enriched for healthy samples (88% and 73% purity) when the third cluster is mostly enriched for T1D samples (81% purity), showing a better separation from our initial clustering base the sole T1D DE signature. Figure 4C provides a visual display of expression values across the ten selected modules with samples sorted by diagnosis and cluster membership. Importantly, we observed a concise expression pattern distinguishing the clusters, in which the diagnosis groups can also be contrasted. This indicates the power of the transcriptomic module analysis in adding robustness to the molecular signature without compromising the specificity of sample separation by diagnosis.

Furthermore, average module expression was calculated per diagnosis and per cluster, per cluster regardless of the diagnosis and per diagnosis regardless of the cluster. We hereafter regard the modules M47 and M29 as exclusive for the cluster “88% healthy”, the modules M2 and M32 as exclusive for the cluster “73% healthy”, and the modules M40, M39, M37, M50, M30 and M44 as exclusive for the cluster “81% T1D”. Similarly, we hereafter consider the modules M2, M32, M47, and M29 as exclusive for healthy samples, and M40, M39, M37, M50, M30 and M44 as exclusive for the cluster T1D samples.

Functional enrichement analyses of T1D-derived gene signatures

To gain more information about the obtained T1D-derived gene signatures, we performed functional enrichement analyses based on the KEGG database.

Collectively, all ten selected modules are enriched for genes up-regulated in T1D belonging to a metabolism-related pathway and down-regulated genes belonging to carbohydrate digestion and absorption pathway (Figure 5A). When we contrast genes belonging to modules over-expressed in T1D samples against genes over-expressed in healthy samples (Figure 5B), we primarily reveal an activation of Th17 cell differentiation pathway in healthy samples not seen in T1D samples. This could seem paradoxical since the accumulation of Th17 have been rather associated to T1D (Fabbri et al., 2019). However, in our study, we look at whole blood transcriptome changes which can be a mirror effect of organ-specific responses. On the contrary, T1D samples show higher expression for genes acting on focal adhesion pathways & PI3K-AKT, two pathways that have been previously linked to insulin metabolism and diabetes (Lebrun et al., 2000).

**Figure 5.**
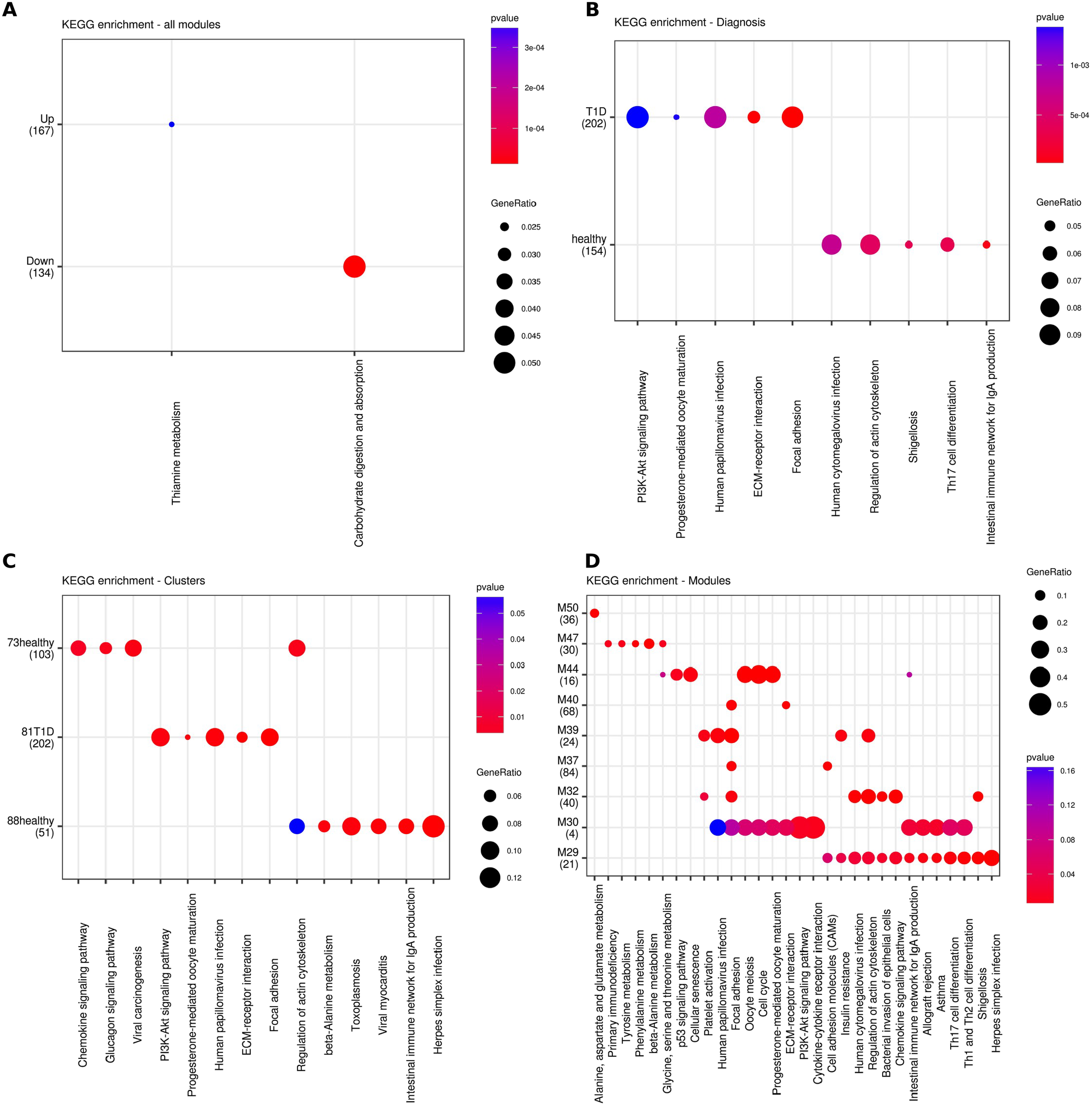
Functional annotation transcriptomic modules. **A)** KEEG enrichment analysis of the ten selected modules merged together and separated by up- and down-regulated. **B)** KEEG enrichment analysis of the modules exclusive for the healthy and T1D samples as delineated in figure 4F. **C)** KEEG enrichment analysis of the modules exclusive for each of clusters, as delineated in figure 4E. **C)** KEEG enrichment analysis of the ten selected modules.

When focusing on groups of modules that are expressed exclusively in clusters (Figure 5C), we observe that the “actin cytoskeleton” is the only pathway over-expressed consistently in the two clusters mostly pure in healthy samples, in line with actin dynamics being important of insulin secretion and pancreatic **↑**-cell biology (Ghitescu et al., 2001; Kalwat and Thurmond, 2013; Lebrun et al., 2000). When zooming in on the enriched pathways in individual modules (Figure 5D), five modules show enrichment of genes acting on Focal adhesion and three for the regulation of actin cytoskeleton, two T1D vs. Healthy discriminative pathways as shown above. Other pathways, mostly related to immune response or metabolism functions, are enriched in one or two modules. Of note, modules M30 & M29 exhibit several enriched pathways due to gene redundancy across pathways.

### Analysis of module expression patterns

Next, we investigated the module expression patterns across the three clusters (Figure 6A-J), to compare their behaviour between the two clusters mostly pure for healthy samples (88% healthy and 73% healthy) and the cluster mostly pure for T1D samples (81% T1D). Four module expression patterns can be described: 1) Modules M30, M44 and M50 show progressive increase in expression across clusters going from the most-prominent healthy (cluster 88% healthy) to the most-prominent T1D (cluster 81% T1D) clusters; 2) Modules M2, M29 and M32 are characterized by greater expression seen in the cluster 73% healthy and lowest expression in cluster 81% T1D; 3) Module M47 alone displays the greatest expression in cluster 88% healthy and lowest expression seen in cluster 73% healthy.

**Figure 6.**
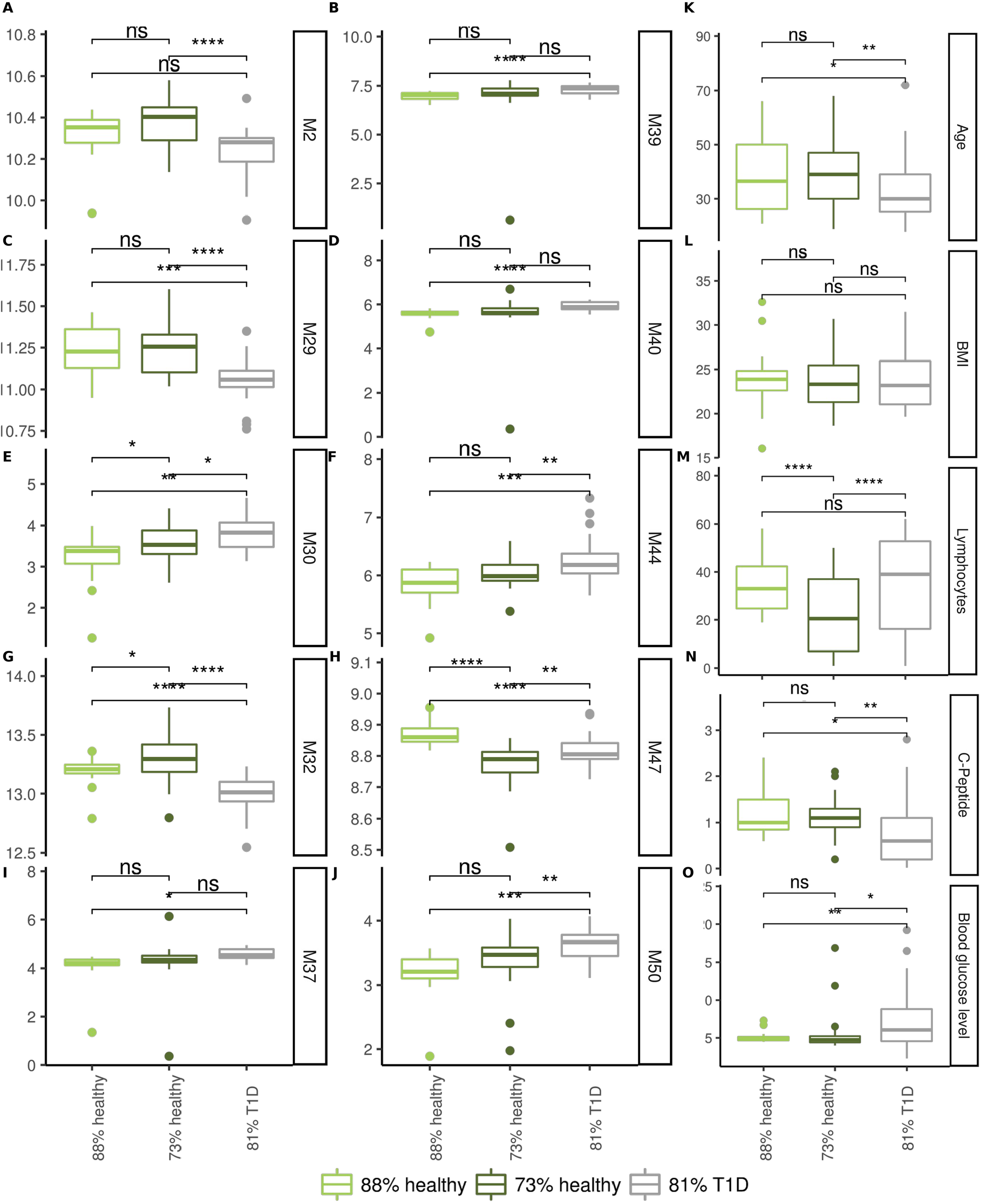
Assessment of modules expressions and cluster composition. **(A-J) Clusters assessment by twosided t-test**. Data is log2 normalized module expression per sample. **(K-O) Values of selected clinical parameters per cluster**. For t-test p-values, ns: p > 0.05, *: p ≤ 0.05, **: p ≤ 0.01, ***: p ≤ 0.001, ****: p ≤ 0.0001

When looking at clinical parameters across clusters (Figure 6K-O), we observe that members of cluster 81% T1D are younger. Cluster 73% healthy display the lowest lymphocyte counts. Not surprisingly, cluster 81% T1D shows the highest blood glucose and the lowest C-Peptide levels. C-peptide level correlates with M2, M29, and M32 expression. Blood glucose level is positively correlated with M30 expression and negatively correlated with M29 and M47 expression. This result is consistent with M30 and M29 modules having KEGG annotations that have been linked to insulin response (i.e. focal adhesion, ECM-receptor interaction, PI3K-akt signalling, insulin resistance, regulation of actin cytoskeleton). Noteworthy these two modules are also well-annotated for immune-response pathways (cytokine-cytokine receptor interaction, chemokine signalling, Th17 cell differentiation, Th1 and Th2 cell differentiation, and infections). Finally, lymphocyte counts correlation negatively with M29 and M32 modules in line with the immune- and infection-related annotations of these modules. Overall, our analysis shows that the constructed modules captured genes involved immune responses or related to amino acid metabolism, insulin signalling, glucagon regulation, functions that have been reported to be associated to diabetes susceptibility (Cui et al., 2009; Fabbri et al., 2019; Ghitescu et al., 2001; Kalwat and Thurmond, 2013; Lebrun et al., 2000; Rajala et al., 2009; Rondas et al., 2012).

### Enrichment of T1D-related genes in modules

We surveyed the functional gene composition of modules by two complementary pathway analysis approaches, namely KEEG pathway enrichment analysis and Ingenuity® Pathway Analysis (Ingenuity Systems, 2013). In total, 954 (mean±sd = 95.4±74.7) genes non-exclusively distributed among the ten selected clusters were considered. A global view on Ingenuity® Pathway Analysis results reveals an enrichment of 68 genes directly implicated on T1D molecular mechanisms (Figure S7 and Table S5). From these, we observed an enrichment of genes belonging to the complement system (C1, C2, and C3), known regulators of immune responses (ICOS, CD200, CD28, and CD34), FACIT collagen family members (COL19A1 and COL6A1), HLA molecules (HLA-DQA1, HLA-DQB2, and HLA-DRB5), integrins (ITGAM, ITGAV, ITGB3), Stat protein family members (STAT3 and STAT6). Two additional enriched proteins (IDE and SIRS1) are implicated in insulin metabolism. We then computed heatmaps aiming at dissecting the module gene expression patterns across samples. Modules M2 and M32 (characteristic of cluster 73% healthy) are remarkably enriched in genes related to “Chemokine signalling” (Figure 5C), including STAT3, CDC42, PLCB2, PXN, PIK3R5, PREX1, CXCR4, and PRKACA. Consistent with observations made from module expressions, gene expression heatmaps show that these genes are down-regulated in cluster 81% T1D and up-regulated to a lesser extent in cluster 88% healthy (Figure 7).

Modules M29 and M47 (characteristic for cluster 88% healthy) are enriched in genes related to “beta-alanine metabolism” and “Intestinal Immune network for IgA production” pathways (see Fig 5C), as shown on respective heatmaps for genes AOC2, AOC3 and ABAT (beta-Alanine metabolism) and auto-immune related genes ICOS, HLA-DQB1, HLA-DRB5 and HLA-DQA1 (Intestinal Immune network for IgA production) (Figure 8). Finally, we focused on modules M40 and M39, characteristic for cluster 81% T1D, enriched for genes related to “Focal adhesion”, including SPP1, VWF, EGF, PDGFRB, FLT1, COL6A1, LAMA2, ITGB3, MYLK, and MYL9. These genes are found over-expressed in all T1D samples regardless of the cluster they belong (Figure 8).

## Conclusion

In this work, we analysed RNAseq transcriptome data of T1D patients and healthy volunteers to identify gene signatures that can best separate sample clusters keeping in mind that heterogeneity of patient transcriptome could spread them in more than one group. We first applied differential expression (DE) analysis to compare whole blood RNA from healthy donors against that of T1D patients.

We produced a first gene list that separates samples from the two compared groups. However, samples did not separate clearly into Healthy vs. T1D clusters, but rather samples spread into three clusters. Functional annotation of the DE genes did not reveal very striking pathways known to underlie T1D molecular mechanisms. Since three clusters were obtained from the DE genes, we hypothesized that either T1D plasticity or sample heterogeneity are playing a role in masking key discriminatory and explanatory genes.

We then used independent component analysis (ICA) to extract additional potential gene signatures from our RNAseq datasets as well as the Molecular Signature Database from the Broad Institute (MSigDB) to identify independently constructed signatures that would be enriched between our samples. We proposed a holistic assessment approach that considers not only the rank produced by gene-set enrichment analysis but also the predictive power and clustering robustness of a given molecular signature. From this approach, we selected 34 significant signatures totalling 3,793 genes. These selected signatures were further combined to construct transcriptomic modules that best capture information to separate samples.

Our results show that jointly combining molecular signatures derived from our expression set (DE and ICA methods) with molecular signatures independently constructed (signatures from MsigDB repository) adds robustness to the analysis without compromising the specificity. This can be seen by the clear separation of the samples according to their diagnosis group when visualizing the module expression and also when analysing the functional annotation of the modules in which a greater number of known T1D-related pathways are represented. Our analysis also reveals high heterogeneity of T1D patient transcriptome data that cannot be well separated from that of healthy volunteers. Instead, T1D patients fall into two main clusters, one T1D-specific cluster and one mixed with healthy volunteers. The constructed transcriptomic show rather clear differential behaviour between clusters showing further that T1D samples do not share clear transcriptomic features as compared to controls.

Together, these results emphasize the challenge to separate T1D patients relative to healthy donors based on their whole blood transcriptomics profiles, with regards to their heterogeneity and despite their common diagnosis. Methodologically, this work outlines that single differential analyses are not able to capture potential immune continuums involved in such a complex pathophysiological process. The stepwise global approach proposed in this work, combining supervised and unsupervised ML-derived approaches, showed the best classification scores.

We hypothesize that T1D patients (i) can have different molecular pathways involved in their pathology; or/and (ii) can display unsynchronized -omics profiles. These findings call for further investigations to identify the molecular pathways involved in T1D patients, and to revise the nosology of autoimmune diseases.

## Contributions

Felipe Leal Valentim (first author, conceived and implemented the solution, wrote the manuscript), Océane Konza (RNAseq data processing and analysis), Alexia Velasquez (GSEA analysis), Ahmed Saadawi (network analysis), Nidhiben Patel (IPA analysis), Roberta Lorenzon (Transimmunom clinical scope), Nicolas Tchitchek (result interpretation, manuscript writing); Mariotti-Ferrandiz (co-supervising author, conception & result interpretation, manuscript writing), David Klatzmann (co-supervising author, conception and immunology scope, result interpretation, manuscript writing), Adrien Six (supervising author, conception and result interpretation, manuscript writing).

## Acknowledgements

Federica Martina (support for conception of framework), Karim El Soufi (support for conception of framework), Michelle Rosenzwajg (support for results interpretation), Fabien Pitoiset (support for results interpretation), Transimmunom consortium.

## Funding

This work was primarily funded by the LabEx Transimmunom (ANR-11-IDEX-0004-02) and RHU iMAP (ANR-16-RHUS-0001) grants, in addition to Sorbonne Université and INSERM recurrent funds. FLV was supported by an ANR grant (ANR-17-ECVD-0005-01).

## Competing interests

The authors declare no competing financial interests.

## Material and Methods

Study participants: type 1 diabetes and healthy volunteers Participants are included in the Clinical Investigation Center at the Pitié-Salpêtrière Hospital. Patients affected by type 1 diabetes are recruited in the departments of Diabetology and Healthy Volunteers (HV) are selected based on internal records in the same hospital.

Participants must be over 18 years old, diagnosed with type 1 diabetes or a healthy subject, and covered by the French healthcare system. Patients are included if they have been diagnosed for type 1 diabetes according to international diagnostic criteria (American Diabetes Association, 2018; Petersmann et al., 2018) for less than 8 years. HV are included, stratified by age in four groups (18-30; 31-40; 41-50; over 50 years old) and matched by sex in each group. A participant, excluded when he/she is undergoing cancer chemotherapy, presents contraindications to donating blood according to (Aigner et al., 2003), is pregnant, is affected by a chronic lifelong viral infection unrelated to the disease, or had an infectious event within the previous month.

The study was approved by the institutional review board of Pitié-Salpêtrière Hospital (ethics committee Ile-De-France 48-15) and done in accordance with the Declaration of Helsinki and good clinical practice. Written informed consent is obtained from all participants before enrolment in the study.

### Clinical data acquisition

The study is conducted by a specialized “Clinical Investigation Center in Biotherapy and Immunology” allowing standardization of the procedure. All data are collected once at the time of clinical evaluation. Peripheral blood samples are collected fasting. Demographic data (sex, age), lifestyle (smoking), disease history, and treatments are collected by physicians based on the clinical document and by querying participants. Body mass index (BMI) is calculated based on the weight and height of the participant, collected on the day of the visit. HbA1C and c-peptide are measured in biochemistry laboratory at the Pitié-Salpêtrière Hospital. IDAA1C score is calculated with daily insulin dose and HbA1c value according to guidelines. All data are collected in anonymized manner and stored (except omics) in a secure e-CRF (Lorenzon et al., 2018). From IDAA1C score we derive remission status; remission when IDAA1C ≤ 9, non-remission when IDAA1C ≥ 9;

### RNA sequencing

Total RNA from whole blood has been extracted following a two-step procedure. First, RNA from blood collected on PAX-Gene tubes has been extracted using Maxwell 16 LEV simplyRNA blood kit (Promega) following manufacturer recommendations and second b-globin, dominant RNA from red blood cells, has been removed using the GLOBINclear kit (Ambion) on extracted RNA. RNA sequencing has been performed from using the TruSeq Stranded mRNA preparation kit (Illumina) on 500 ng b-globin depleted RNA with a RNA Integrity Number > 8 (measured on Bioanalyzer following manufacturer recommendations), and then sequenced following a pairend 2×75 bp protocol on NextSeq 500 or HiSeq 4000 (Illumina) at LIGAN Equipex (Lille, France).

### Quantification of Protein expression

Starting from the raw fastq files containing RNA-seq paired reads for each sample, quality control reports were generated using FastQC (Andrews, 2010). Additionally, possible contamination was assessed by screening the reads against several databases using FastQ Screen (Wingett, 2017). Overall, the primary FastQC quality reports of the data show satisfactory patterns, and no contamination was detected by FastQ Screen. MultiQC (Ewels et al., 2016) was used to generate summary reports of these results (Figure S1)

The reads of each sample were then aligned against the Human genome using default parameters of hisat2 for paired reads of forward-stranded library. The genome index database was built over the Ensembl Human genome (Yates et al., 2016) version GRCh37. Next, featureCounts (Liao et al., 2014) parameters was also adjusted for analysis of paired reads of forward-stranded library at gene level. The gene annotation used here was also obtained from Ensembl version GRCh37. The read counts for each sample was merged in a final table and analysed with R (Hornik, 2018) scripts developed specifically for this dataset. Only protein coding genes located on autosomal chromosomes were further considered.

### Putative signatures

Two distinct approaches were used to obtain putative signatures; namely, count-based differential expression (Anders et al., 2013) and independent component analysis (Hyvärinen, 2013). From the myriad of methods available for count-based differential expression, “DESeq2” (Love et al., 2014) was used in combination with standard tests; namely “roc” (standard area-under-the-curve classifier), “t” (student’s t-test) and “wilcox” (Wilcoxon rank sum test). These latest tests were used as implemented in the R package Seurat (Satija et al., 2015).

First, the raw read counts were normalized following specific recommendations for each of the methods. The DESeq2 analysis requires the definition of the experimental design formula. Here, the gender of the sample donors and the sample batches were used as co-variates whilst the diagnosis was used as phenotype. Second, a DESeq2 p-value threshold was set to 5.10-5, bellow which the gene is included in the putative signature. For the standard tests, the normalization scheme started from the normalized counts from the DESeq2 object. Then, the batch effect was explicitly treated by applying removeBatchEffect function from R package limma (Ritchie et al., 2015). The p-value thresholds were also set to 1.10-5 for both wilcox and t tests, values below which the gene is included in the putative signature. As for the roc test, a threshold of 0.65 was adjusted for the predictive power of each gene, values above which the gene is included in the putative signature. The signatures obtained from each of these differential expression methods are grouped into the DE signatures. Moreover, the union of the DE genes obtained from the different methods was considered to generate the T1D DE signature.

Independent component analysis (ICA) were also performed on the same batch-effect-treated dataset as the one used by the standard tests. Default ICA parameters were then used as implemented by (Pham et al., 2014) (icaAlpha=1.0, maxIteration=1000, tol= 1.10-5, icaIteration=100, icaCorr=0.95, icaThreshold=3). The gene signatures generated by ICA framework are referred to as ICA signatures.

Independent collection of annotated gene sets

In addition to DE and ICA signatures, whole collections of gene sets were obtained from the Molecular Signatures database (MsigDB) (Liberzon et al., 2015). Specifically, the C7 (immunologic signatures) and the C2 (curated gene sets) were downloaded from the MsigDB repository. These are referred to as C2 and C7 signatures.

### Signature assessment

All four groups of signatures (i.e. DE, ICA, C2, and C7) were considered. Additionally, one set of 1000 randomly simulated gene signatures was created. For this, a signature length l was randomly picked from a simulated normal distribution obtained from the mean and standard deviation of the original signature lengths. This strategy was implemented with the MCMCglmm R Package (Hadfield, 2010), which allows to truncate the distribution, in such a way only numbers from a defined range are sampled. The length limit was adjusted as [50,500]. Once having set the length l, l protein coding genes located on autosomal chromosomes were randomly selected to create a random signature. The Random signature group was thus created repeating this procedure 1000 times.

To account for heterogeneity of the data, 100 sub-datasets were created by bootstrapping 50% of the samples 100x. Each of the signatures in all groups were then evaluated by three independent groups of analyses, i.e. predictive power, robustness and gene-set enrichment analysis. Analyses were also performed on the bootstrapped sub-datasets.

Firstly, the accuracy of predictions obtained by predictive models was used as an estimate of the predictive power of each signature. These predictive models were constructed based solely on the expression of the genes in a given signature combined with the sample phenotype. Two predictive modelling approaches were used: Random Forest (Díaz-Uriarte and Alvarez de Andrés, 2006) and Support Vector Machine (Vanitha et al., 2015). Random Forest (RF) was implemented with R package randomForest (Breiman, 2001) and Support Vector Machine (SVM) with R package caret (Kuhn, 2008). For SVM model default parameters were used, and for RF ntree=500 was used. In both cases, the same strategy was used as follow: the 50% of the bootstrapped samples were used to train the predictive model, whilst the 50% of samples left out from the bootstraps were used to compute the accuracy of the predictions. The model is trained with expressions of genes in a given signature as variables whilst the phenotype of the samples (diagnosis) are considered the predictors. The overall accuracy is then computed from the confusion matrix of predictions against the original sample diagnosis.

Secondly, the Calinski-Harabasz (CH) index (Caliñski and Harabasz, 1974) was used as an estimate of the clustering Robustness. This CH index was computed based solely on the expression of the genes in a given signature combined with the sample labels. Here, the sample labels were set as either 1) the Diagnosis (Robustness for Diagnosis) or, 2) the cluster in which the sample falls into when hierarchical clustering is applied to the gene set (Robustness for hClust). The hierarchical clustering was implemented with R package stats (Hornik, 2018) by applying the algorithm: a distance matrix is initially constructed from gene expression (function dist, method=“Euclidean”); then hierarchical clustering is applied to construct the hierarchical tree (function hclust, method=“complete”); and finally the cluster labels are obtained by cutting the tree into four clusters (function cutree, k=4). The CH index was implemented with R package clues (Chang et al., 2010) with default parameters using either the Diagnosis labels or the cluster labels.

Thirdly, gene-set enrichment analysis was used to assess the enrichment of each signature. For that, an algorithm for preranked gene set enrichment analysis using cumulative statistic calculation was used as implemented in the R package fgsea (Sergushichev, 2016).The preranked gene set enrichment analysis requires as input a gene ranking list, where each gene is ranked according to its differential expression between the two groups (healthy donors and T1D). To create this ranking list, the t test stat computed between the mean expression in each group was used. After creating the list, fgsea on all signatures is executed. Then, signatures are sorted by enrichment p-values, where the lowest p-value is the first in the rank (GSEA RNK).

p-values were computed in a similar manner for all evaluated metrics. All 100 bootstrapped sub-datasets were used; thus 100 values for each evaluated metric are obtained to assess each signature in the main four groups (DE, ICA, C2, and C7). Similarly, 100 values for each evaluated metric are also obtained for each of the 1000 signatures in the Random group. Therefore, the t-test was performed between the 100 values of the evaluated metric for a given signature against the diagnostic values obtained for all the Random signatures. To select the top signatures, the following criteria were used: a p-value threshold was set, below which the signature is further considered; then 10 signatures with the most prominent values for the metric are taken as top 10. A similar p-value < 5.10-3 threshold was used for all metrics. The average between the values obtained from the 100 bootstrapped sub-datasets is used for the construction of the histograms only. The statistic results are shown in Table S2.

### Cluster-based expression fingerprinting

In a first step, our method we use gap statistics to estimate the optimum number of gene clusters (k) for each molecular signature; then k gene clusters are computed. In the second step, we calculate a distance matrix between each gene cluster using binary distance. From this binary distance matrix, a distance tree is created. Here, each node of the distance tree represents one of the clusters gene cluster obtained from selected molecular signatures (Figure S11). The ‘modules’ are then created by cutting neighbouring nodes in the binary distance tree. Finally, the average expression of genes in each ‘module’ is computed for each sample.

The R package multiClust (Lawlor et al., 2016) was used to compute the optimal number of gene clusters for each molecular signatures using gap statistics. To obtain the binary distance matrix among clusters of all signatures, R method dist (method=“binary”) (Hornik, 2018) was used. From distance matrix, hClust (method = “ward.D”) was used to compute the distance tree. Finally, R method cutree (k=70) was used to retrieve the modules from the distance tree.

The modular expression is given by averaging the expression all genes in each module per sample. The modular expression was first assessed by chi-square test of expression against diagnostic. Additionally, t-test between modular expression of healthy samples and T1D samples was performed. Finally, a random forest model was constructed with expressions of all modules, then variable importance analysis was performed to rank the modules accordingly.

### Data availability

Transcriptomic RNA-seq data have deposited on the Gene Expression Omnibus database and are available via the accession number GSE123658.

**Figure 7A.**
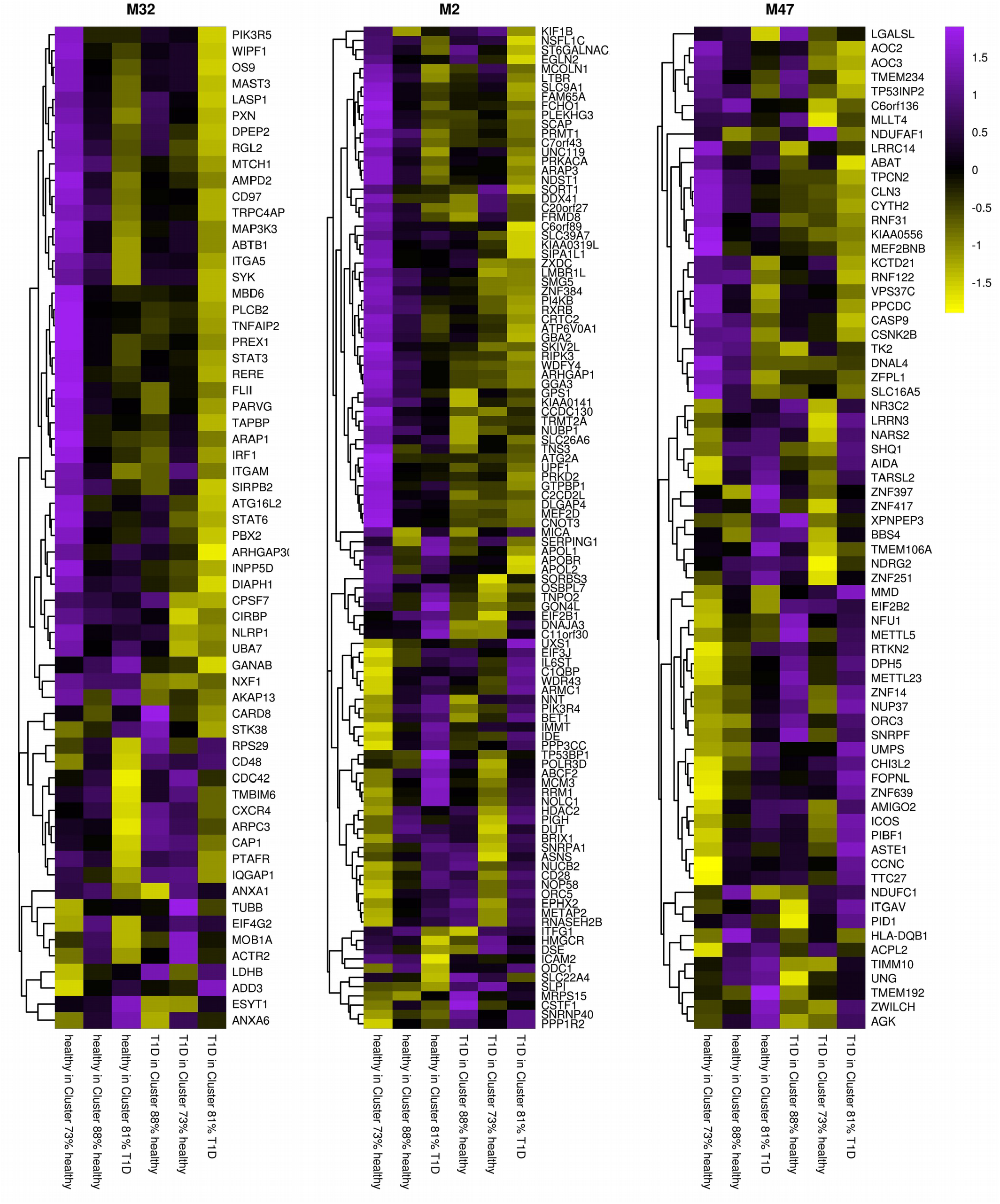
Gene composition of selected transcriptomic modules. A**-C)** Heatmaps with gene expressions per group for the selected modules (M32, M2 and M47). Each group is composed by combining one of the two diagnostic groups (healthy of T1D) with one of the three clusters (88% healthy, 73% healthy and 81%T1D). The expression represents the expression of the single gene over the samples of each group.

**Figure 7B.**
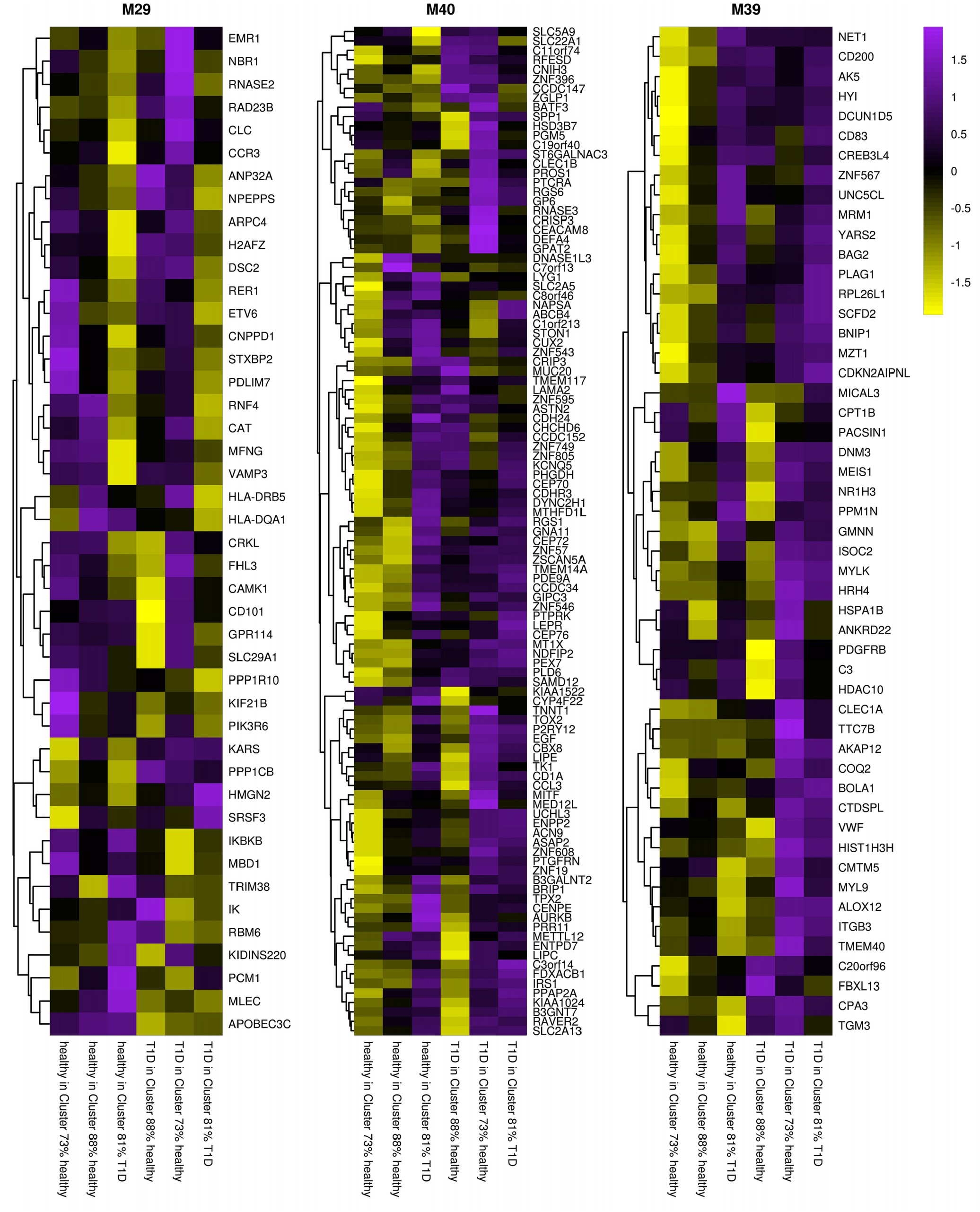
Gene composition of selected transcriptomic modules. A**-C)** Heatmaps with gene expressions per group for the selected modules (M29, M40 and M39). Each group is composed by combining one of the two diagnostic groups (healthy of T1D) with one of the three clusters (88% healthy, 73% healthy and 81%T1D). The expression represents the expression of the single gene over the samples of each group.

## Notes

### Competing Interest Statement

The authors have declared no competing interest.

