## Supplemental Figure for "Transcriptomic module fingerprint reveals heterogeneity of whole blood transcriptome in type 1 diabetic patients"

### Heterogeneity Aware Evaluation of Gene signatures on whole blood from type 1 diabetic patients

Felipe Leal Valentim (first author, conceived and implemented the solution),Sophie Harris (manager, scenario thinking), Encarnita (facilitator, co-supervisor, results interpretation), David (conceived experiments, definition of immunology scope), Adrien Six (supervising author, conceived the ideas), Ahmed.saadawi (network analysis), Federica (support for conception of framework), Karim (support for conception of framework), Michelle (support for results interpretation), Roberta (definition of clinical scope), Fabien (support for results interpretation), Rna-extraction (missing name???), Alexia (conceived ideas for GSEA analysis, analysis of feasibility of GSEA for transimmunom data), Nidhiben Patel (conceived ideas for IPA analysis, analysis of feasibility of IPA for transimmunom data), LIGAN (Veronique) - Service providers, not co-authors

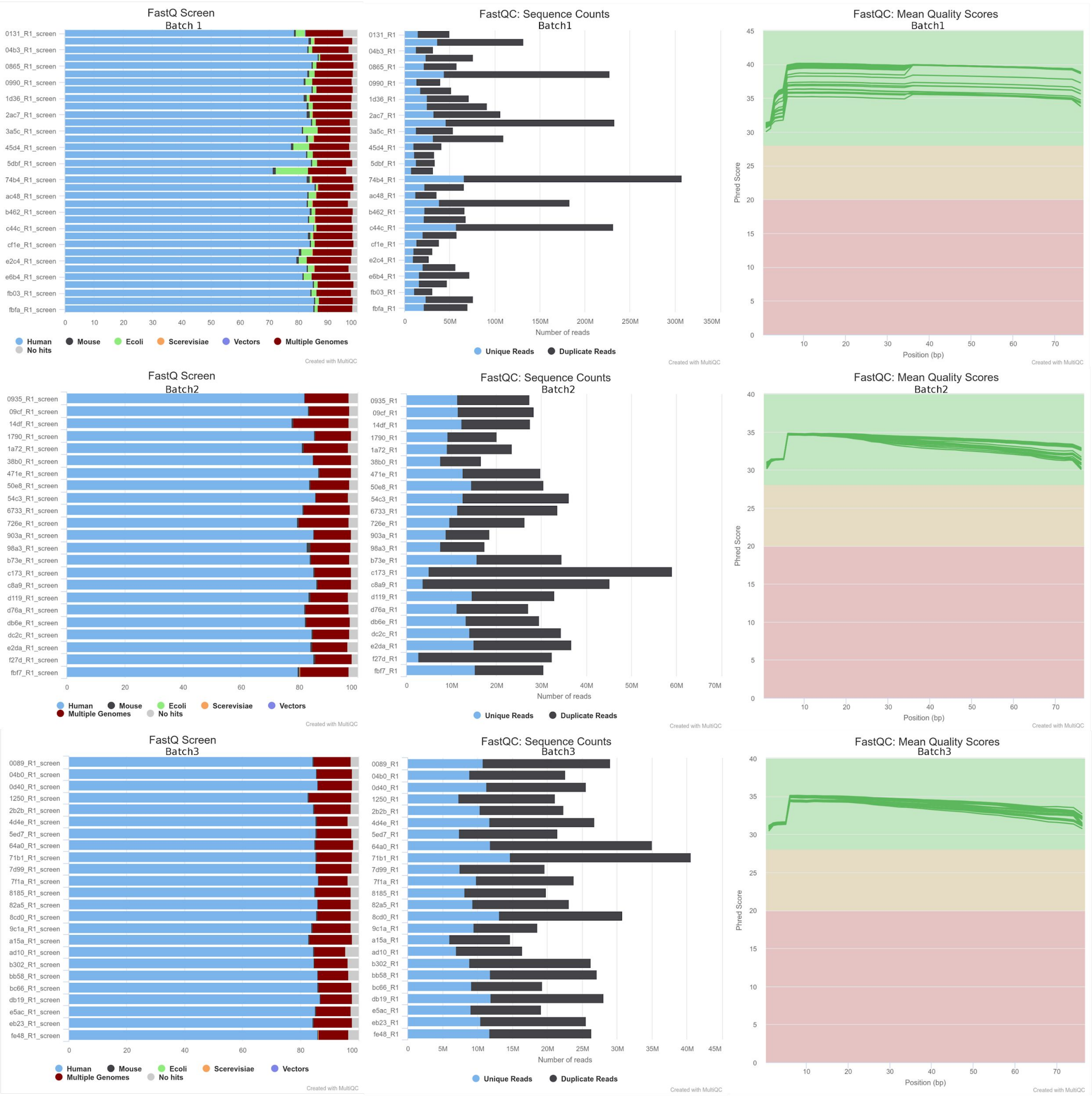
FastQ Screen
Batch 1
0131_R1_screen
04b3_R1_screen
0865_R1_screen
0990_R1_screen
1d36_R1_screen
2ac7_R1_screen
3a5c_R1_screen
45d4_R1_screen
5dbf_R1_screen
74b4_R1_screen
ac48_R1_screen
b462_R1_screen
c44c_R1_screen
cf1e_R1_screen
e2c4_R1_screen
e6b4_R1_screen
fb03_R1_screen
fbfa_R1_screen
0 10 20 30 40 50 60 70 80 90 100
Human Mouse Ecoli Scerevisiae Vectors Multiple Genomes No hits
Created with MultiQC
FastQC: Sequence Counts
Batch1
0131_R1
04b3_R1
0865_R1
0990_R1
1d36_R1
2ac7_R1
3a5c_R1
45d4_R1
5dbf_R1
74b4_R1
ac48_R1
b462_R1
c44c_R1
cf1e_R1
e2c4_R1
e6b4_R1
fb03_R1
fbfa_R1
0 50M 100M 150M 200M 250M 300M 350M
Number of reads
Unique Reads Duplicate Reads
Created with MultiQC
FastQC: Mean Quality Scores
Batch1
Phred Score
0 5 10 15 20 25 30 35 40 45
10 20 30 40 50 60 70
Position (bp)
Created with MultiQC
FastQ Screen
Batch2
0935_R1_screen
09cf_R1_screen
14df_R1_screen
1790_R1_screen
1a72_R1_screen
38b0_R1_screen
471e_R1_screen
50e8_R1_screen
54c3_R1_screen
6733_R1_screen
726e_R1_screen
903a_R1_screen
98a3_R1_screen
b73e_R1_screen
c173_R1_screen
c8a9_R1_screen
d119_R1_screen
d76a_R1_screen
db6e_R1_screen
dc2c_R1_screen
e2da_R1_screen
f27d_R1_screen
fbf7_R1_screen
0 20 40 60 80 100
Human Mouse Ecoli Scerevisiae Vectors Multiple Genomes No hits
Created with MultiQC
FastQC: Sequence Counts
Batch2
0935_R1
09cf_R1
14df_R1
1790_R1
1a72_R1
38b0_R1
471e_R1
50e8_R1
54c3_R1
6733_R1
726e_R1
903a_R1
98a3_R1
b73e_R1
c173_R1
c8a9_R1
d119_R1
d76a_R1
db6e_R1
dc2c_R1
e2da_R1
f27d_R1
fbf7_R1
0 10M 20M 30M 40M 50M 60M 70M
Number of reads
Unique Reads Duplicate Reads
Created with MultiQC
FastQC: Mean Quality Scores
Batch2
Phred Score
0 5 10 15 20 25 30 35 40
10 20 30 40 50 60 70
Position (bp)
Created with MultiQC
FastQ Screen
Batch3
0089_R1_screen
04b0_R1_screen
0d40_R1_screen
1250_R1_screen
2b2b_R1_screen
4d4e_R1_screen
5ed7_R1_screen
64a0_R1_screen
71b1_R1_screen
7d99_R1_screen
7f1a_R1_screen
8185_R1_screen
82a5_R1_screen
8cd0_R1_screen
9c1a_R1_screen
a15a_R1_screen
ad10_R1_screen
b302_R1_screen
bb58_R1_screen
bc66_R1_screen
db19_R1_screen
e5ac_R1_screen
eb23_R1_screen
fe48_R1_screen
0 20 40 60 80 100
Human Mouse Ecoli Scerevisiae Vectors Multiple Genomes No hits
Created with MultiQC
FastQC: Sequence Counts
Batch3
0089_R1
04b0_R1
0d40_R1
1250_R1
2b2b_R1
4d4e_R1
5ed7_R1
64a0_R1
71b1_R1
7d99_R1
7f1a_R1
8185_R1
82a5_R1
8cd0_R1
9c1a_R1
a15a_R1
ad10_R1
b302_R1
bb58_R1
bc66_R1
db19_R1
e5ac_R1
eb23_R1
fe48_R1
0 5M 10M 15M 20M 25M 30M 35M 40M 45M
Number of reads
Unique Reads Duplicate Reads
Created with MultiQC
FastQC: Mean Quality Scores
Batch3
Phred Score
0 5 10 15 20 25 30 35 40
10 20 30 40 50 60 70
Position (bp)
Created with MultiQC

### S1 Figure

**Summary of quality control and contamination screen for all samples per batch.** Contamination screen of the sequences against a set of databases; sequence counts for each sample and respective estimate of duplicate read counts; the mean quality value across each base position in the read.

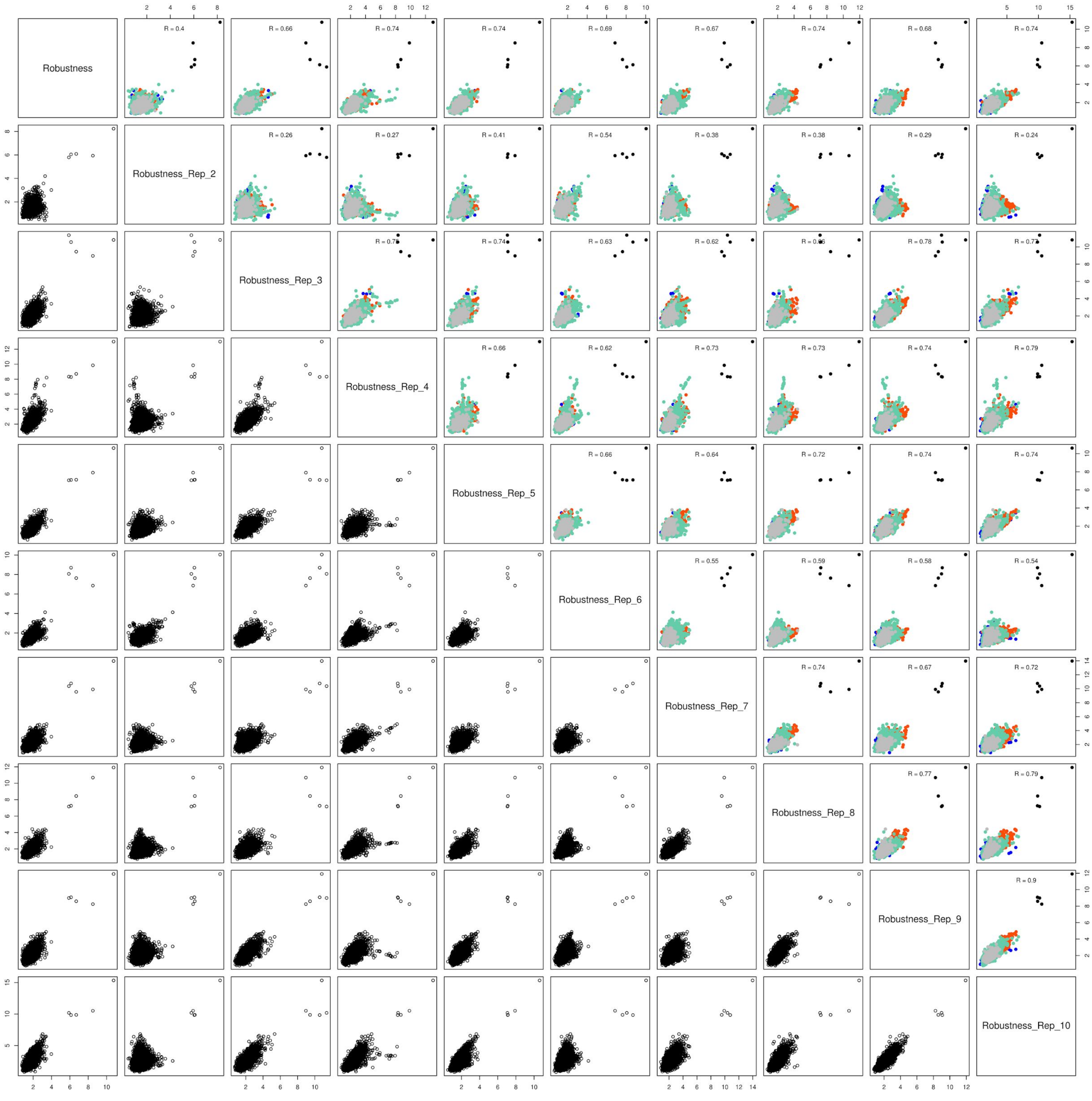

Robustness
Robustness_Rep_2
Robustness_Rep_3
Robustness_Rep_4
Robustness_Rep_5
Robustness_Rep_6
Robustness_Rep_7
Robustness_Rep_8
Robustness_Rep_9
Robustness_Rep_10
R = 0.4
R = 0.66
R = 0.74
R = 0.74
R = 0.69
R = 0.67
R = 0.74
R = 0.68
R = 0.74
R = 0.26
R = 0.27
R = 0.41
R = 0.54
R = 0.38
R = 0.38
R = 0.29
R = 0.24
R = 0.74
R = 0.63
R = 0.62
R = 0.78
R = 0.66
R = 0.62
R = 0.73
R = 0.73
R = 0.74
R = 0.79
R = 0.66
R = 0.64
R = 0.72
R = 0.74
R = 0.74
R = 0.55
R = 0.59
R = 0.58
R = 0.54
R = 0.74
R = 0.67
R = 0.72
R = 0.77
R = 0.79
R = 0.9

### S2 Figure

**Correlation matrix for Robustness (over Diagnosis groups) computed from different bootstrapped sub-datasets.** Only results over the 10 first boostrapped set-datasets are  plotted duo to graphical constraints

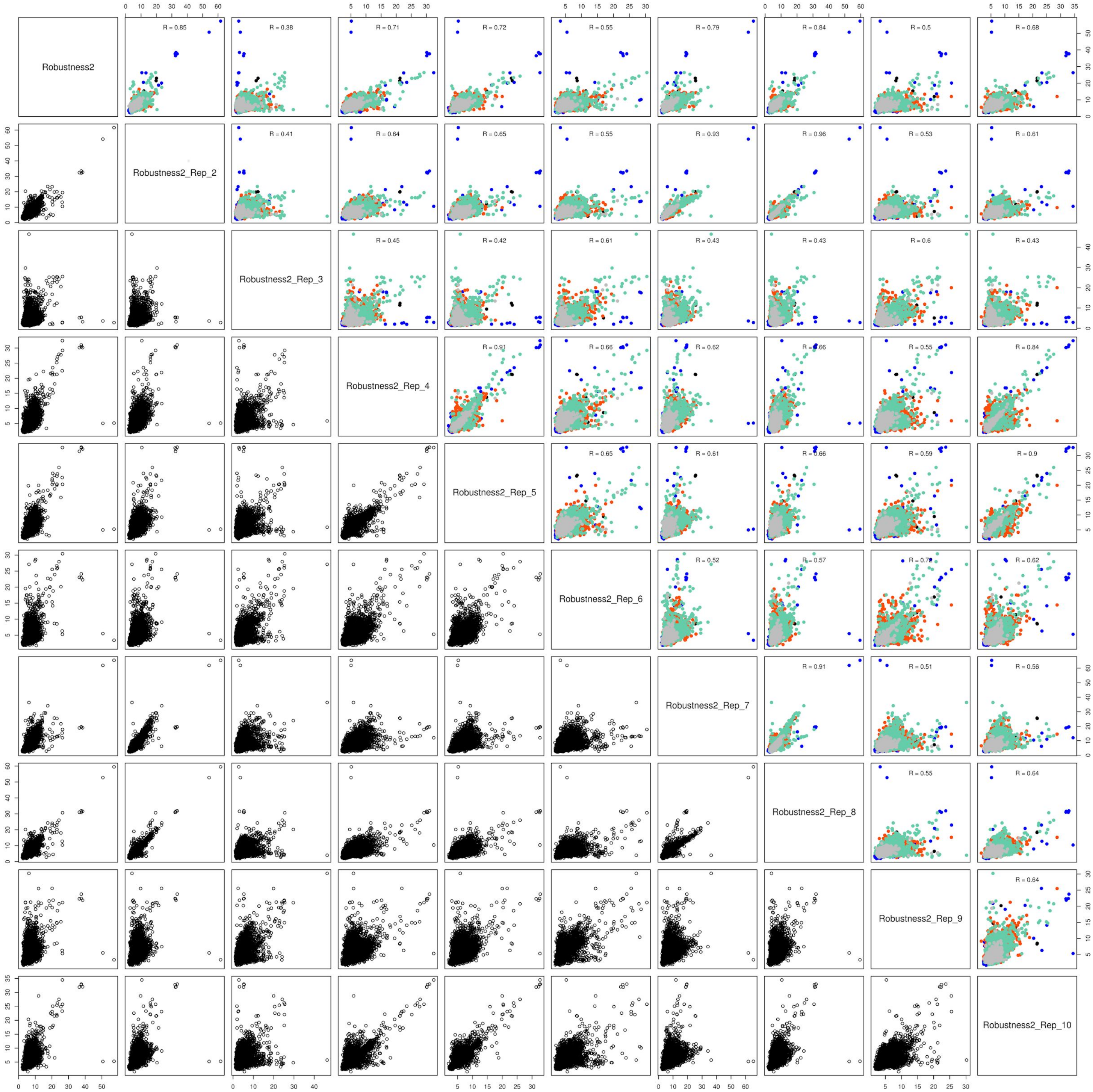

Robustness2
Robustness2_Rep_2
Robustness2_Rep_3
Robustness2_Rep_4
Robustness2_Rep_5
Robustness2_Rep_6
Robustness2_Rep_7
Robustness2_Rep_8
Robustness2_Rep_9
Robustness2_Rep_10
R = 0.85
R = 0.38
R = 0.71
R = 0.72
R = 0.55
R = 0.79
R = 0.84
R = 0.5
R = 0.68
R = 0.41
R = 0.64
R = 0.65
R = 0.55
R = 0.93
R = 0.96
R = 0.53
R = 0.61
R = 0.45
R = 0.42
R = 0.61
R = 0.43
R = 0.43
R = 0.6
R = 0.43
R = 0.91
R = 0.66
R = 0.62
R = 0.66
R = 0.55
R = 0.84
R = 0.65
R = 0.61
R = 0.66
R = 0.59
R = 0.9
R = 0.52
R = 0.57
R = 0.72
R = 0.62
R = 0.91
R = 0.51
R = 0.56
R = 0.55
R = 0.64
R = 0.64

### S3 Figure

**Correlation matrix for Robustness (over hClust labels) computed from different bootstrapped sub-datasets.** Only results over the 10 first boostrapped set-datasets are  plotted duo to graphical constraints.

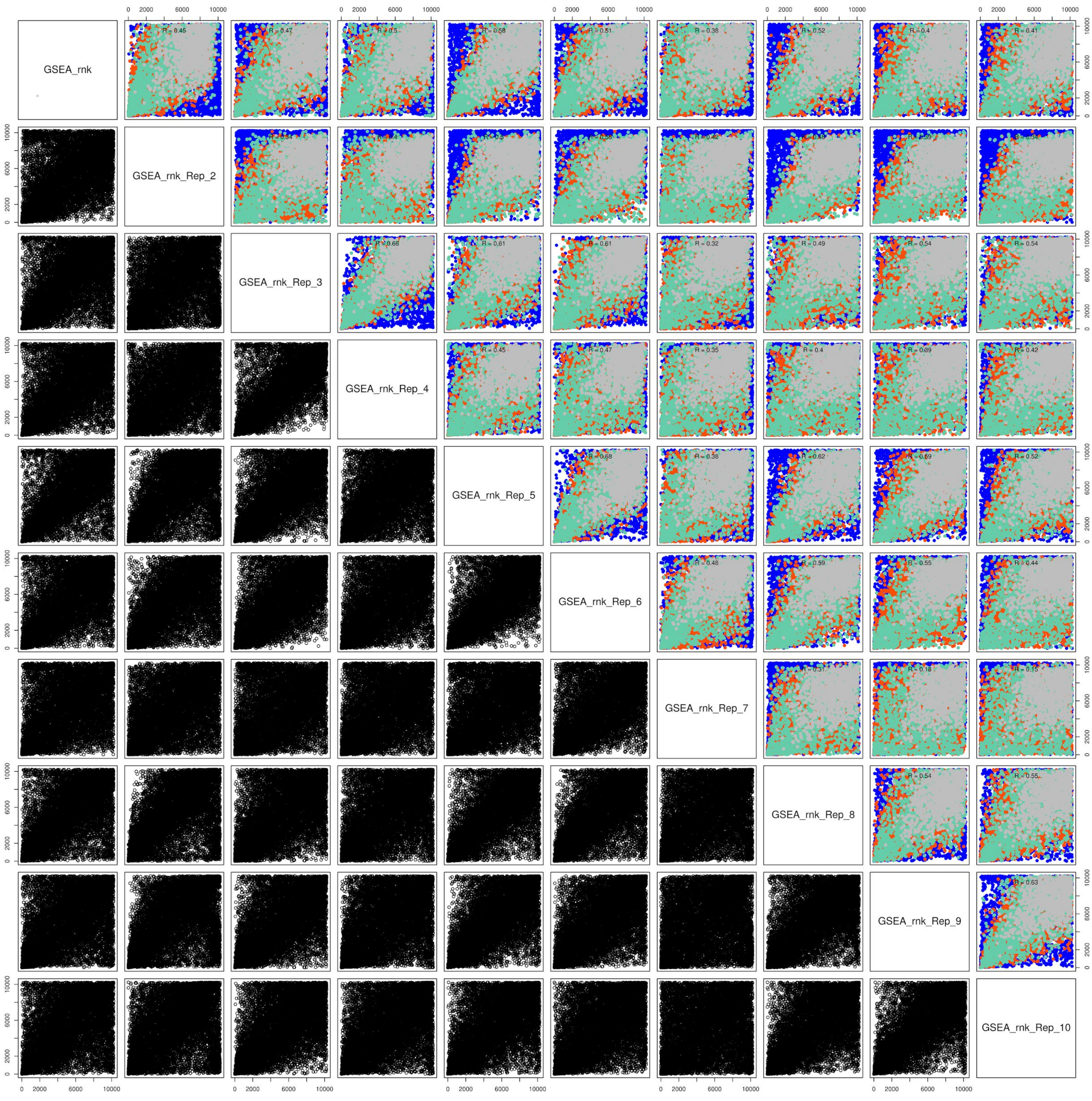
GSEA_rnk
GSEA_rnk_Rep_2
GSEA_rnk_Rep_3
GSEA_rnk_Rep_4
GSEA_rnk_Rep_5
GSEA_rnk_Rep_6
GSEA_rnk_Rep_7
GSEA_rnk_Rep_8
GSEA_rnk_Rep_9
GSEA_rnk_Rep_10

### S4 Figure

**Correlation matrix for signature rank (from GSEA analysis) computed from different bootstrapped sub-datasets.** Only results over the 10 first boostrapped set-datasets are  plotted duo to graphical constraints.

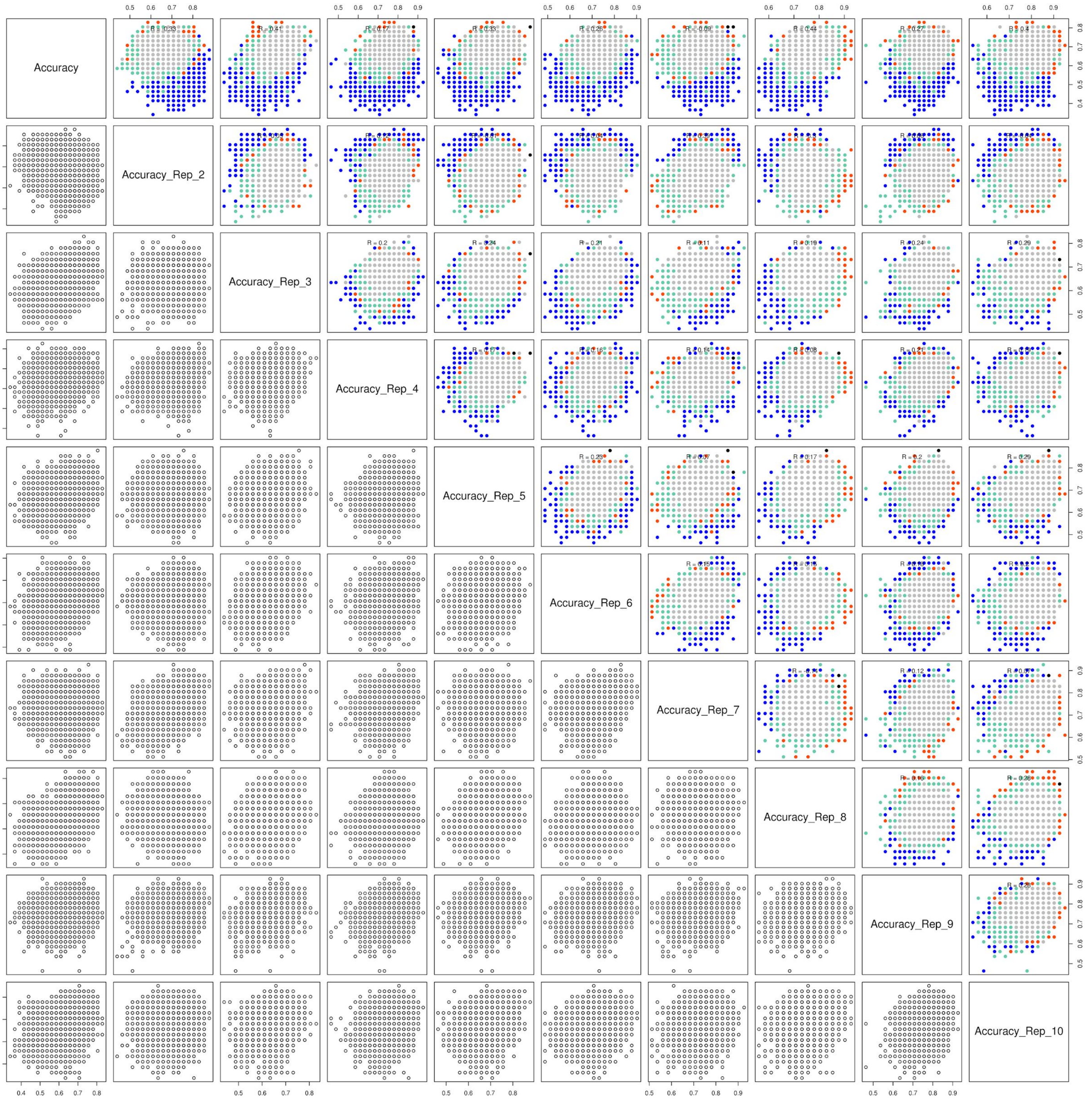
Accuracy
Accuracy_Rep_2
Accuracy_Rep_3
Accuracy_Rep_4
Accuracy_Rep_5
Accuracy_Rep_6
Accuracy_Rep_7
Accuracy_Rep_8
Accuracy_Rep_9
Accuracy_Rep_10

### S5 Figure

**Correlation matrix for Accuracy (from RF predictions) computed from different bootstrapped sub-datasets.** Only results over the 10 first boostrapped set-datasets are plotted duo to graphical constraints.

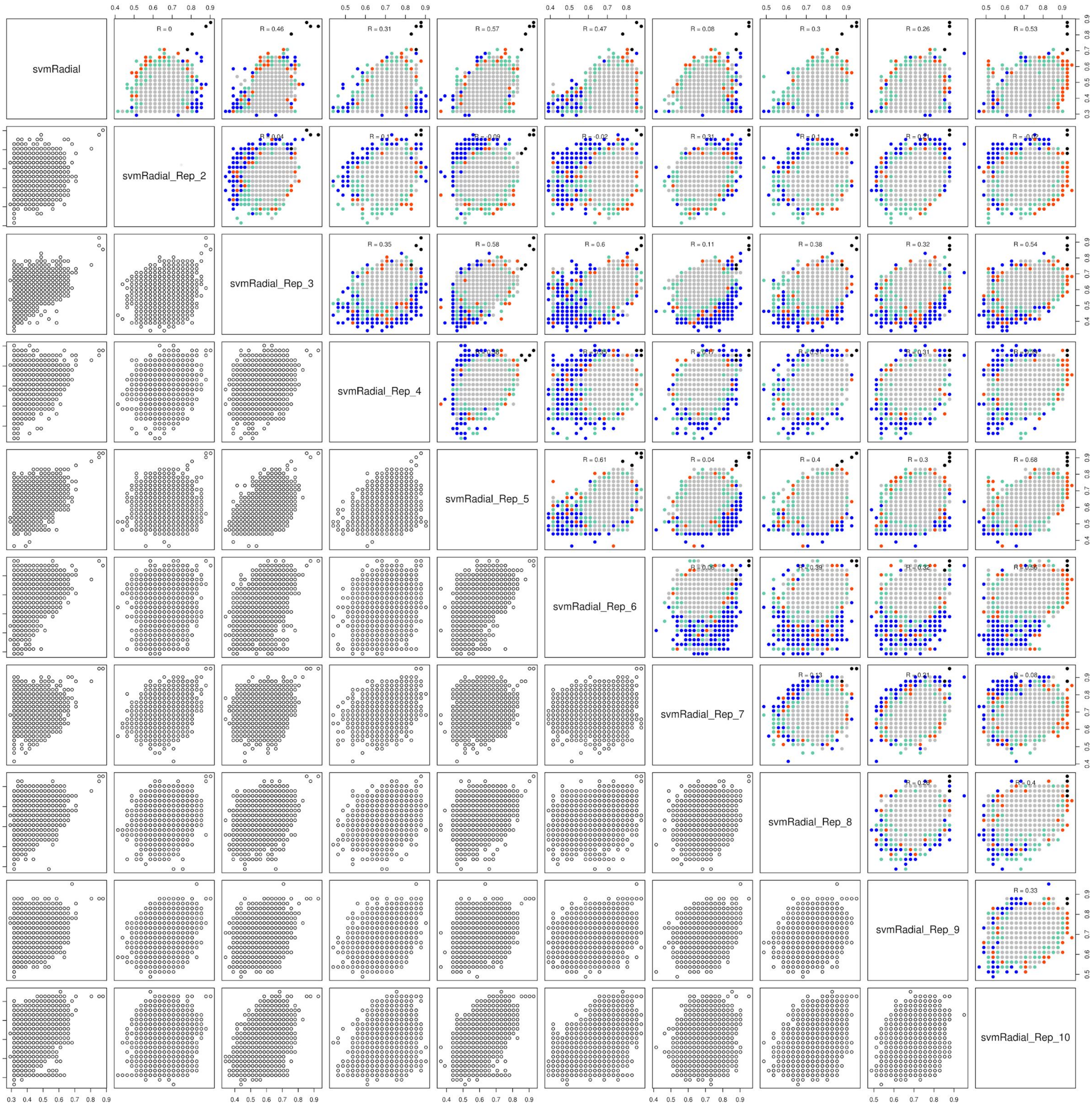
svmRadial
svmRadial_Rep_2
svmRadial_Rep_3
svmRadial_Rep_4
svmRadial_Rep_5
svmRadial_Rep_6
svmRadial_Rep_7
svmRadial_Rep_8
svmRadial_Rep_9
svmRadial_Rep_10
R = 0
R = 0.46
R = 0.31
R = 0.57
R = 0.47
R = 0.08
R = 0.3
R = 0.26
R = 0.53
R = 0.35
R = 0.58
R = 0.6
R = 0.11
R = 0.38
R = 0.32
R = 0.54
R = 0.61
R = 0.04
R = 0.4
R = 0.3
R = 0.68
R = 0.33

### S6 Figure

**Correlation matrix for Accuracy (from SVM predictions) computed from different bootstrapped sub-datasets.** Only results over the 10 first boostrapped set-datasets are plotted duo to graphical constraints.

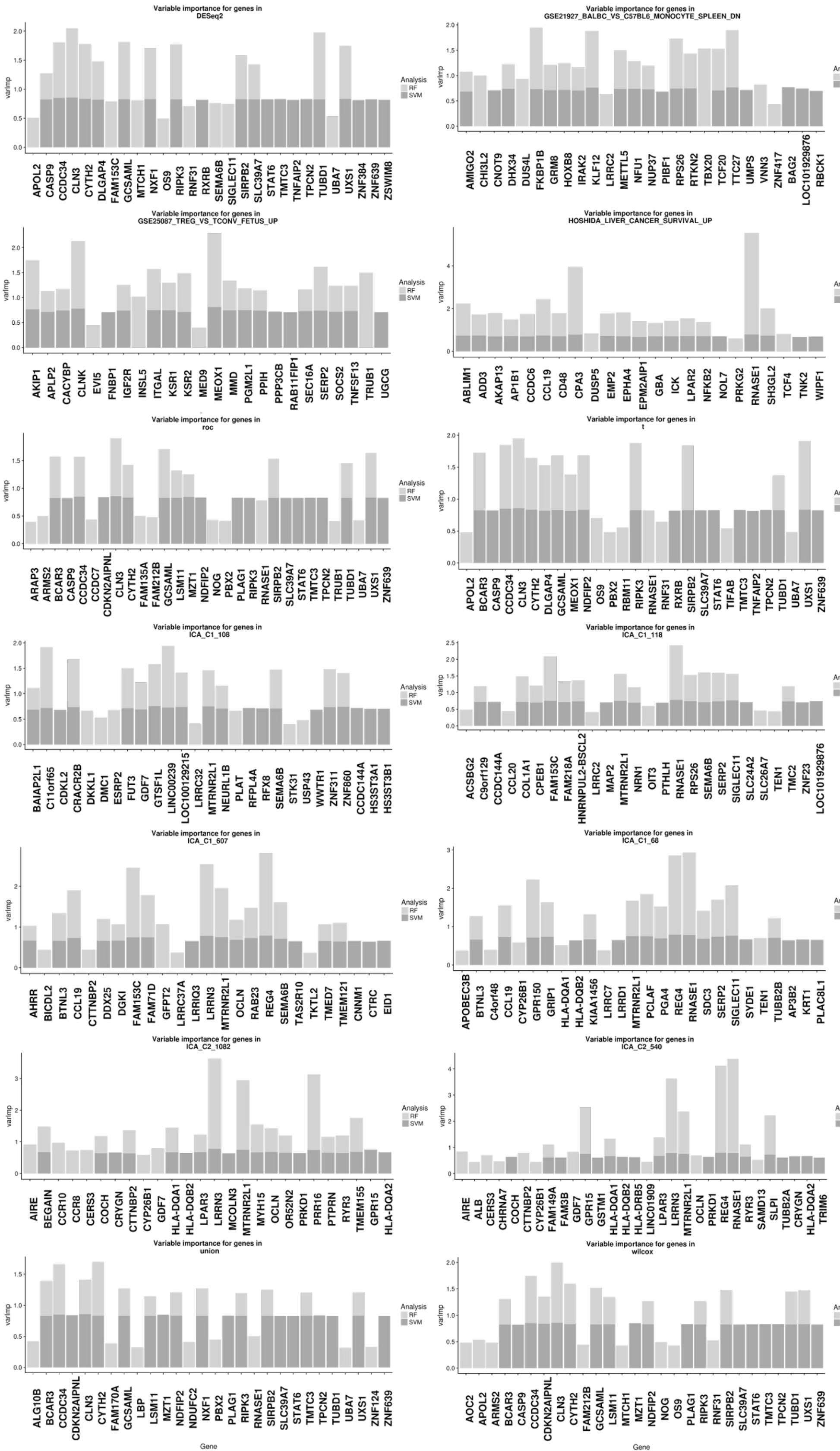
Variable importance for genes in DESeq2
Variable importance for genes in GSE21927_BALBC_VS_C57BL6_MONOCYTE_SPLEEN_DN
Variable importance for genes in GSE25087_TREG_VS_TCONV_FETUS_UP
Variable importance for genes in HOSHIDA_LIVER_CANCER_SURVIVAL_UP
Variable importance for genes in roc
Variable importance for genes in t
Variable importance for genes in ICA_C1_108
Variable importance for genes in ICA_C1_118
Variable importance for genes in ICA_C1_607
Variable importance for genes in ICA_C1_68
Variable importance for genes in ICA_C2_1082
Variable importance for genes in ICA_C2_540
Variable importance for genes in union
Variable importance for genes in wilcox
Analysis
RF
SVM
varImp
Gene

### S7 Figure

**Variable importance analysis of signatures enriched by predictive power assessment.** Variable importance analysis was performed on enriched signatures by both Support Vector Machine and Random Forest models. The top 20 important genes from both models are combined to create the panels, which each shows results for signature indicated by panel title.

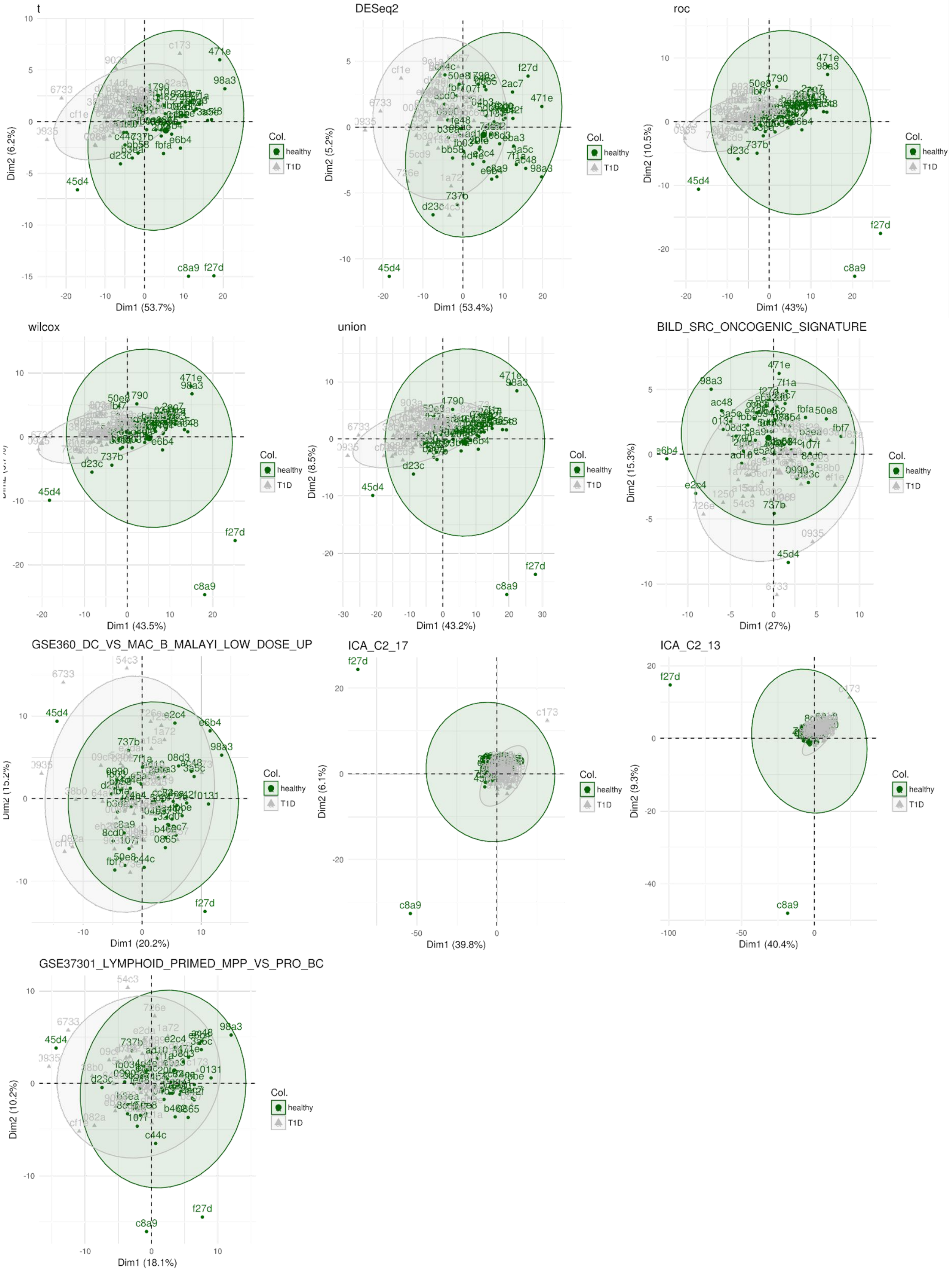
t
Dim1 (53.7%)
Dim2 (6.2%)
DESeq2
Dim1 (53.4%)
Dim2 (5.2%)
roc
Dim1 (43%)
Dim2 (10.5%)
wilcox
Dim1 (43.5%)
union
Dim1 (43.2%)
Dim2 (8.5%)
BILD_SRC_ONCOGENIC_SIGNATURE
Dim1 (27%)
Dim2 (15.3%)
GSE360_DC_VS_MAC_B_MALAYI_LOW_DOSE_UP
Dim1 (20.2%)
ICA_C2_17
Dim1 (39.8%)
Dim2 (6.1%)
ICA_C2_13
Dim1 (40.4%)
Dim2 (9.3%)
GSE37301_LYMPHOID_PRIMED_MPP_VS_PRO_BC
Dim1 (18.1%)
Dim2 (10.2%)
Col.
healthy
T1D

### S8 Figure

**Analysis clustering robustness from diagnosis labels.** Principal component analysis computed with genes of each enriched signature. Samples are colored green or grey, representing respectively healthy or T1D samples.

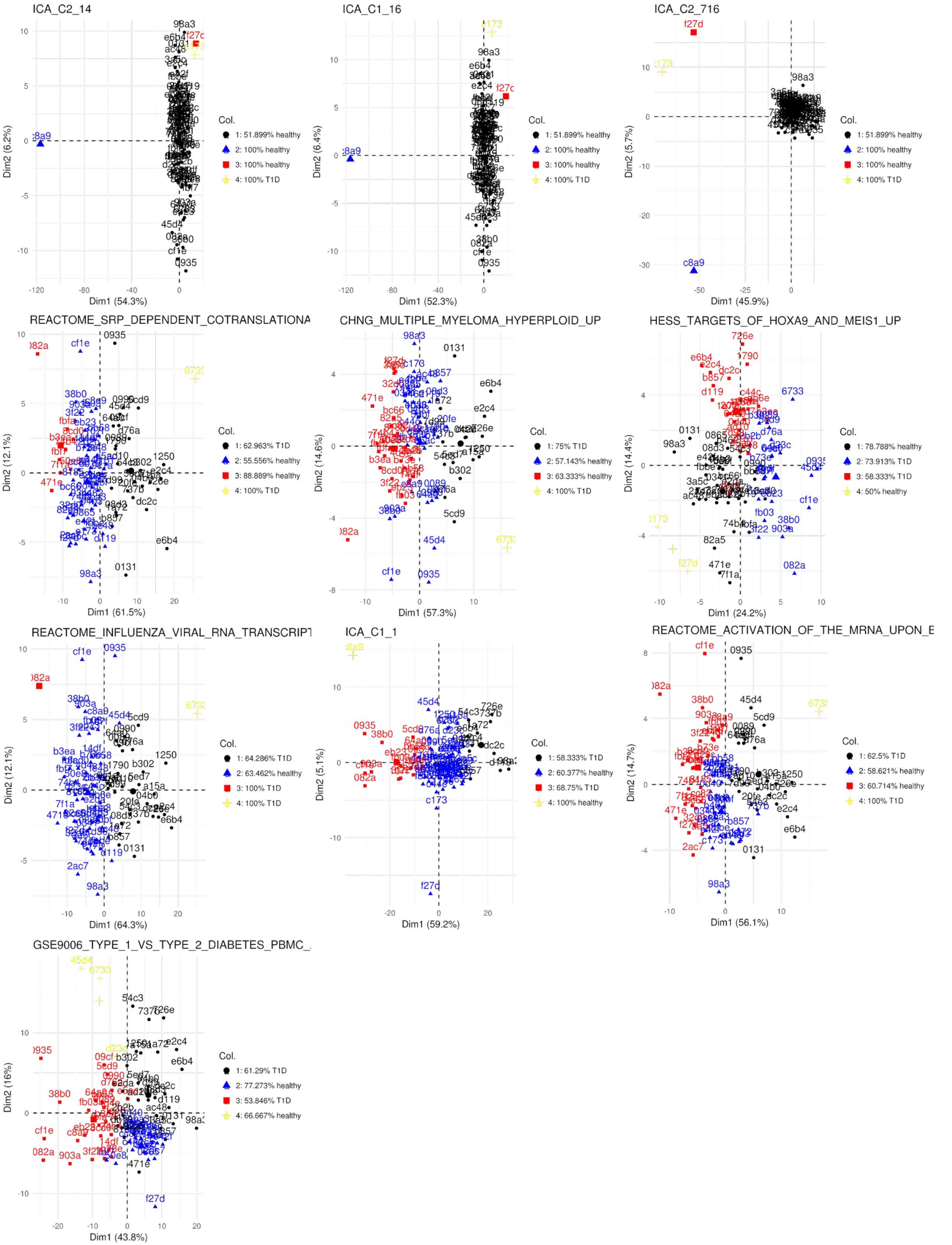
ICA_C2_14
Dim1 (54.3%)
Dim2 (6.2%)
Col.
1: 51.899% healthy
2: 100% healthy
3: 100% healthy
4: 100% T1D
ICA_C1_16
Dim1 (52.3%)
Dim2 (6.4%)
Col.
1: 51.899% healthy
2: 100% healthy
3: 100% healthy
4: 100% T1D
ICA_C2_716
Dim1 (45.9%)
Dim2 (5.7%)
Col.
1: 51.899% healthy
2: 100% healthy
3: 100% healthy
4: 100% T1D
REACTOME_SRP_DEPENDENT_COTRANSLATIONA
Dim1 (61.5%)
Dim2 (12.1%)
Col.
1: 62.963% T1D
2: 55.556% healthy
3: 88.889% healthy
4: 100% T1D
CHNG_MULTIPLE_MYELOMA_HYPERPLOID_UP
Dim1 (57.3%)
Dim2 (14.6%)
Col.
1: 75% T1D
2: 57.143% healthy
3: 63.333% healthy
4: 100% T1D
HESS_TARGETS_OF_HOXA9_AND_MEIS1_UP
Dim1 (24.2%)
Dim2 (14.4%)
Col.
1: 78.788% healthy
2: 73.913% T1D
3: 58.333% T1D
4: 50% healthy
REACTOME_INFLUENZA_VIRAL_RNA_TRANSCRIPT
Dim1 (64.3%)
Dim2 (12.1%)
Col.
1: 64.286% T1D
2: 63.462% healthy
3: 100% T1D
4: 100% T1D
ICA_C1_1
Dim1 (59.2%)
Dim2 (5.1%)
Col.
1: 58.333% T1D
2: 60.377% healthy
3: 68.75% T1D
4: 100% healthy
REACTOME_ACTIVATION_OF_THE_MRNA_UPON_E
Dim1 (56.1%)
Dim2 (14.7%)
Col.
1: 62.5% T1D
2: 58.621% healthy
3: 60.714% healthy
4: 100% T1D
GSE9006_TYPE_1_VS_TYPE_2_DIABETES_PBMC_
Dim1 (43.8%)
Dim2 (16%)
Col.
1: 61.29% T1D
2: 77.273% healthy
3: 53.846% T1D
4: 66.667% healthy

### S9 Figure

**Analysis of robustness of clusters computed with hierarchical clustering.** Principal component analysis computed with genes of each enriched signature. K=4 sample clusters were computed with genes in each signature. The clusters are highlighted by the colors, and corresponding cluster purites are indicated in the legend.

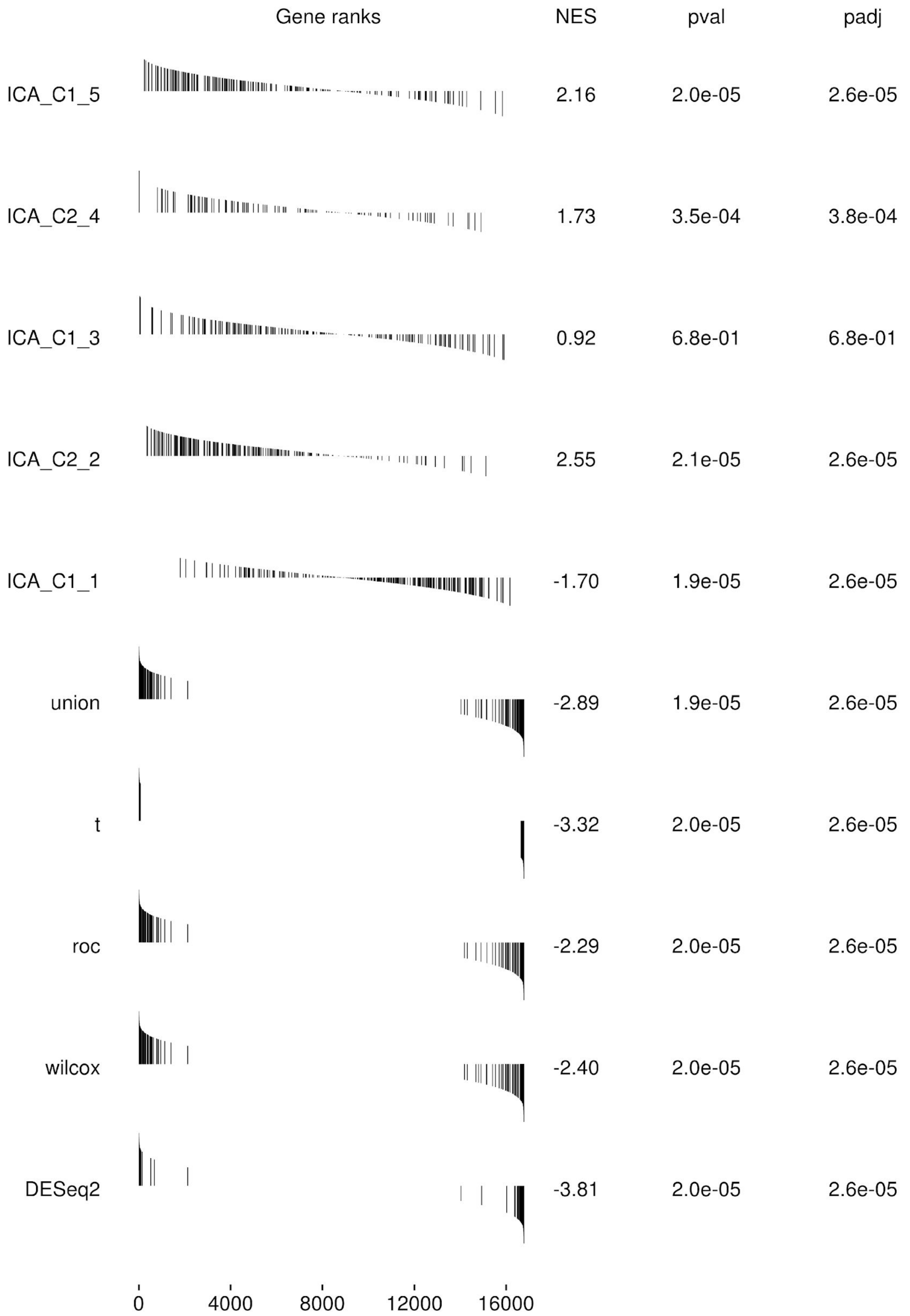
Gene ranks
NES
pval
padj
ICA_C1_5
2.16
2.0e-05
2.6e-05
ICA_C2_4
1.73
3.5e-04
3.8e-04
ICA_C1_3
0.92
6.8e-01
6.8e-01
ICA_C2_2
2.55
2.1e-05
2.6e-05
ICA_C1_1
-1.70
1.9e-05
2.6e-05
union
-2.89
1.9e-05
2.6e-05
t
-3.32
2.0e-05
2.6e-05
roc
-2.29
2.0e-05
2.6e-05
wilcox
-2.40
2.0e-05
2.6e-05
DESeq2
-3.81
2.0e-05
2.6e-05
0
4000
8000
12000
16000

### S10 Figure

**Summary of gene-set-enrichment analysis.** Enrichment plot, normalized enrichment score, p-value (pval) and adjusted p-value (padj) for top 10 signatures enriched by GSEA.

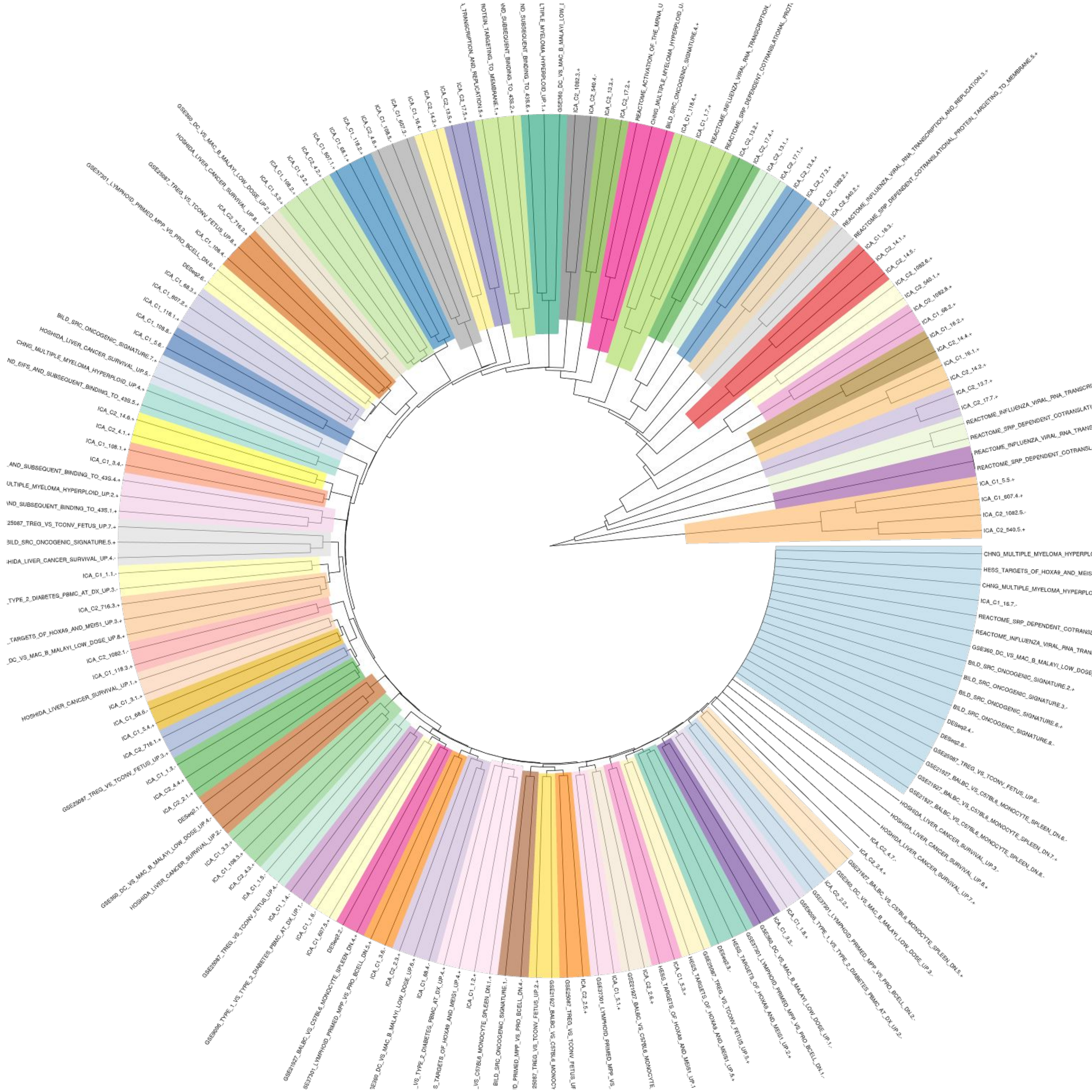
ICA_C1_5.5.+
ICA_C1_607.4.+
ICA_C2_1082.5.-
ICA_C2_540.5.+
ICA_C1_16.7.-
DESeq2.4.-
DESeq2.8.-
BILD_SRC_ONCOGENIC_SIGNATURE.5.+
ICA_C1_1.1.-
ICA_C2_716.3.+
ICA_C2_1082.1.-
ICA_C1_118.3.+
HOSHIDA_LIVER_CANCER_SURVIVAL_UP.1.+
ICA_C1_3.4.-
ICA_C1_108.1.+
ICA_C2_4.1.+
ICA_C2_14.6.+

### S11 Figure

**Phylogenetic tree from distance matrix among all sub-clusters from the enriched moluecular signatures.** The clusters are grouped in modules (delineated by different colors). The label following each cluster indicates the name of the molecular signature from which it belongs follow by a numerical identifier.

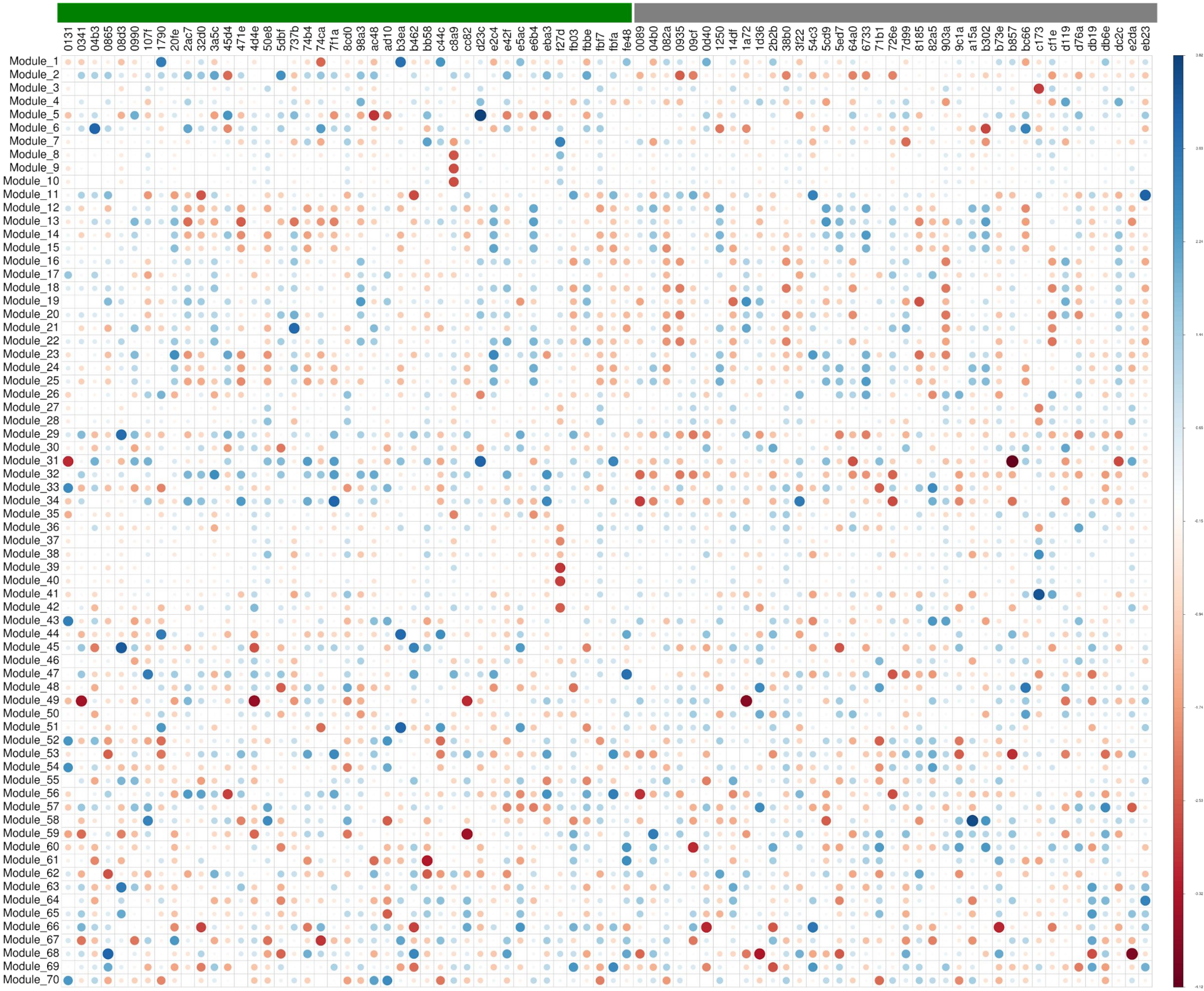

0131
0341
04b3
0865
08d3
0990
107f
1790
20fe
2ac7
32d0
3a5c
45d4
471e
4d4e
50e8
5dbf
737b
74b4
74ca
7f1a
8cd0
98a3
ac48
ad10
b3ea
b462
bb58
c44c
c8a9
cc82
d23c
e2c4
e42f
e5ac
e6b4
eba3
f27d
fb03
fbbe
fbf7
fbfa
fe48
0089
04b0
082a
0935
09cf
0d40
1250
14df
1a72
1d36
2b2b
38b0
3f22
54c3
5cd9
5ed7
64a0
6733
71b1
726e
7d99
8185
82a5
903a
9c1a
a15a
b302
b73e
b857
bc66
c173
cf1e
d119
d76a
db19
db6e
dc2c
e2da
eb23
Module_1
Module_2
Module_3
Module_4
Module_5
Module_6
Module_7
Module_8
Module_9
Module_10
Module_11
Module_12
Module_13
Module_14
Module_15
Module_16
Module_17
Module_18
Module_19
Module_20
Module_21
Module_22
Module_23
Module_24
Module_25
Module_26
Module_27
Module_28
Module_29
Module_30
Module_31
Module_32
Module_33
Module_34
Module_35
Module_36
Module_37
Module_38
Module_39
Module_40
Module_41
Module_42
Module_43
Module_44
Module_45
Module_46
Module_47
Module_48
Module_49
Module_50
Module_51
Module_52
Module_53
Module_54
Module_55
Module_56
Module_57
Module_58
Module_59
Module_60
Module_61
Module_62
Module_63
Module_64
Module_65
Module_66
Module_67
Module_68
Module_69
Module_70

### S12 Figure

**Module expressions per sample.** The module expression is given by averaging the expression of the genes that compose it. The figure show the module expression per sample. The module expression is averaged by cluster of diagnosis depending on the analysis.

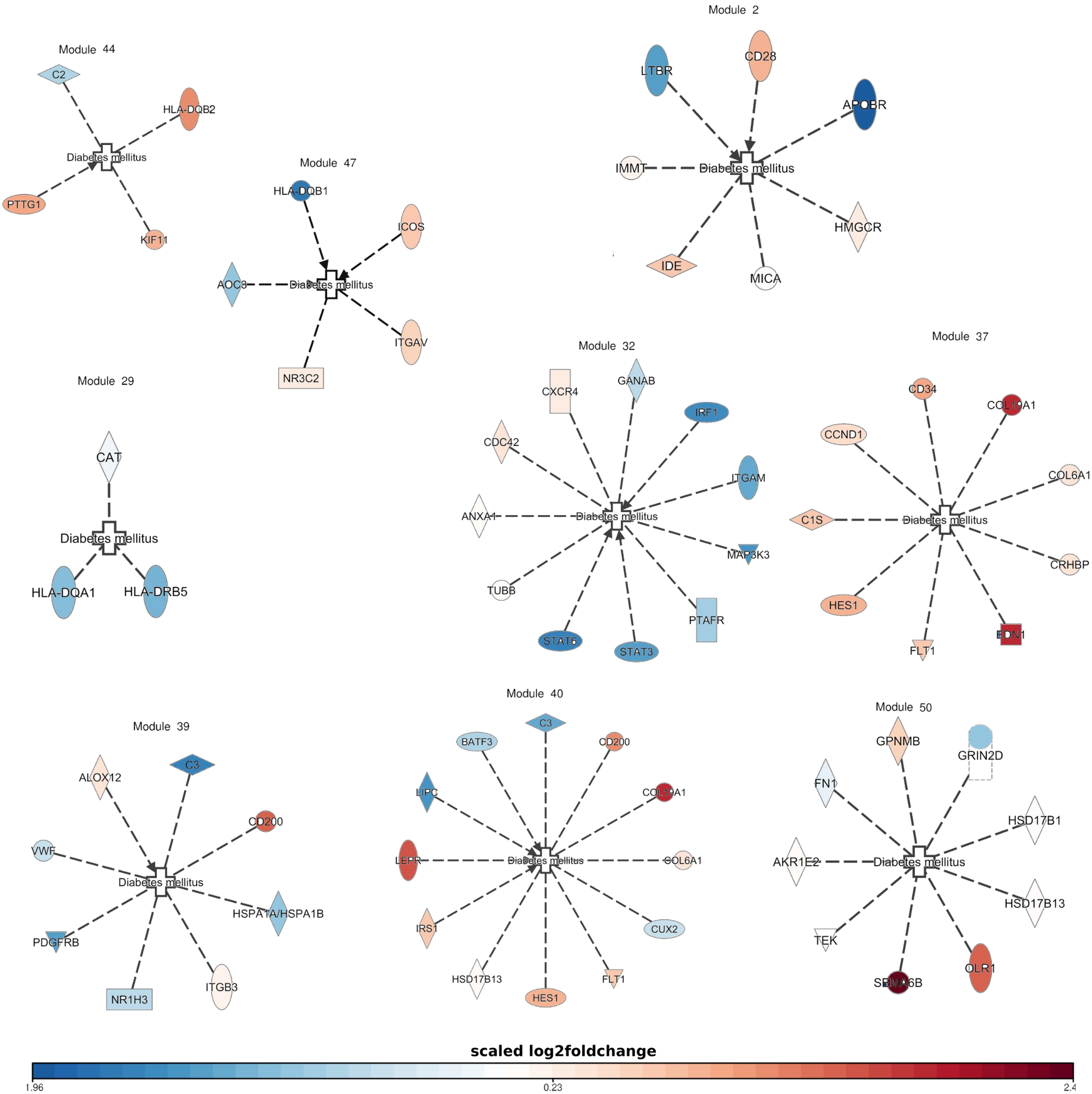
Module 44
C2
HLA-DQB2
Diabetes mellitus
PTTG1
KIF11
Module 47
HLA-DQB1
ICOS
AOC3
Diabetes mellitus
ITGAV
NR3C2
Module 2
LTBR
CD28
APOBR
IMMT
Diabetes mellitus
HMGCR
IDE
MICA
Module 29
CAT
Diabetes mellitus
HLA-DQA1
HLA-DRB5
Module 32
CXCR4
GANAB
IRF1
CDC42
ITGAM
ANXA1
Diabetes mellitus
MAP3K3
TUBB
PTAFR
STAT6
STAT3
Module 37
CD34
COL19A1
CCND1
COL6A1
C1S
Diabetes mellitus
CRHBP
HES1
EDN1
FLT1
Module 39
ALOX12
C3
CD200
VWF
Diabetes mellitus
HSPA1A/HSPA1B
PDGFRB
NR1H3
ITGB3
Module 40
C3
BATF3
CD200
LIPC
COL19A1
LEPR
Diabetes mellitus
COL6A1
IRS1
CUX2
HSD17B13
HES1
FLT1
Module 50
GPNMB
GRIN2D
FN1
HSD17B1
AKR1E2
Diabetes mellitus
HSD17B13
TEK
SEMA6B
OLR1
scaled log2foldchange
1.96
0.23
2.4

### S13 Figure

**Documented T1D-related genes within selected modules.** The genes in the modules where compare against the T1D-related genes documented by the Ingenuity Knowledge Base (Quiagen®).
